# Meiotic recombination modulates the structure and dynamics of the synaptonemal complex during *C. elegans* meiosis

**DOI:** 10.1101/110064

**Authors:** Divya Pattabiraman, Baptiste Roelens, Alexander Woglar, Anne M. Villeneuve

**Author notes:** These authors contributed equally to this work (Divya Pattabiraman, Baptiste Roelens, Alexander Woglar). Corresponding Author: Anne M. Villeneuve Department of Developmental Biology 279 Campus Dr. B300 Beckman Center Stanford University School of Medicine Stanford, CA 94305.

## Abstract

During meiotic prophase, a structure called the synaptonemal complex (SC) assembles at the interface between aligned pairs of homologous chromosomes, and crossover recombination events occur between their DNA molecules. Here we investigate the inter-relationships between these two hallmark features of the meiotic program in the nematode *C. elegans*, revealing dynamic properties of the SC that are modulated by recombination. We demonstrate that the SC incorporates new subunits and switches from a more highly dynamic/labile state to a more stable state as germ cells progress through the pachytene stage of meiotic prophase. We further show that the more dynamic state of the SC is prolonged in mutants where meiotic recombination is impaired. Moreover, in meiotic mutants where recombination intermediates are present in limiting numbers, SC central region subunits become preferentially stabilized on the subset of chromosome pairs that harbor a site where pro-crossover factors COSA-1 and MutS_γ_ are concentrated. Polo-like kinase PLK-2 becomes preferentially localized to the SCs of chromosome pairs harboring recombination sites prior to the enrichment of SC central region proteins on such chromosomes, and PLK-2 is required for this enrichment to occur. Further, late pachytene nuclei in a *plk-2* mutant exhibit the more highly dynamic SC state. Together our data demonstrate that crossover recombination events elicit chromosome-autonomous stabilizing effects on the SC and implicate PLK-2 in this process. We discuss how this recombination-triggered modulation of SC state might contribute to regulatory mechanisms that operate during meiosis to ensure the formation of crossovers while at the same time limiting their numbers.

## Author Summary

Reliable chromosome inheritance during sexual reproduction depends on the formation of temporary connections between homologous chromosomes that enable them to segregate toward opposite spindle poles at the meiosis I division. These connections are established during an extended meiotic prophase characterized by two prominent features: a highly-ordered structure called the synaptonemal complex (SC) that assembles at the interface between aligned pairs of chromosomes, and crossover recombination events between their DNA molecules that are completed in the context of the SC. In the current work, we investigate the inter-relationships between these two hallmark features of the meiotic program in the nematode *C. elegans*. Our work reveals the *C. elegans* SC as a much more dynamic structure than is suggested by its highly-ordered appearance in EM images, and further demonstrates that the SC switches from a more highly dynamic/labile state to a more stable state as germ cells progress through meiotic prophase. Moreover, we show that formation of crossover recombination intermediates can trigger stabilization of the SC in a chromosome autonomous manner. We speculate that recombination-triggered SC stabilization may provide a means for germ cells to monitor whether chromosome pairs have acquired the prerequisite crossover intermediate needed to ensure correct homolog segregation.

## Introduction

Sexual reproduction depends on the specialized cell division program of meiosis, which allows diploid organisms to form haploid gametes. Reduction in chromosome number from the diploid to the haploid state occurs during the first meiotic division, when homologous chromosomes segregate to opposite poles of the meiosis I spindle. Reliable segregation of homologs in turn depends on creation of temporary attachments between the homologs during an extended prophase that precedes the meiotic divisions. In most organisms, meiotic prophase is characterized by two hallmark features: 1) assembly of a highly-ordered structure known as the synaptonemal complex (SC) between the aligned homologs, and 2) formation of crossover (CO) recombination events between their DNA molecules (1).

The occurrence of crossing over between homologs during meiosis was recognized more the 100 years ago (2), long before DNA was identified as the genetic material. We now know that COs are the products of a meiosis-specific recombination program, consisting of i) the controlled introduction of DNA double strand breaks (DSBs) by the transesterase SPO-11 (3–5), ii) engagement of the homologous chromosome as a template for recombinational repair (6), and iii) formation and resolution of intermediates at a subset of recombination sites in a manner that yields CO products. These inter-homolog COs, in conjunction with sister chromatid cohesion, form the basis of connections that allow the homologs to orient and segregate toward opposite spindle poles at meiosis I. Meiotic CO formation is promoted by conserved meiosis-specific factors that include the MutS_γ_ (MSH4-MSH5) complex (7–10), and in animals, the cyclin-related protein COSA-1/CNTD1 (11, 12). Further, meiotic recombination is tightly regulated in a manner that limits that number of COs formed, yet simultaneously guarantees that every homolog pair receives at least one CO.

The SC has also long been recognized as a canonical feature of the meiotic program (13, 14). In EM images, and more recently through use of super-resolution fluorescence microscopy, the SC is observed as a tripartite structure assembled at the interface between pairwise aligned homologous chromosomes (15–18). Axial structures that form along the length of each homolog comprise the two lateral elements of the SC, paralleling each other at an approximate distance of 100-200 nm. These lateral elements are linked together by the SC central region, which contains an ordered array of transverse filaments that spans the distance between the two axes, resulting in a zipper- or railroad track-like appearance of the SC in EM images. Meiotic chromosome axes in most organisms contain meiosis-specific cohesin complexes and/or meiotic HORMA-domain-containing proteins (however, the numbers of parologous complexes present in different organisms is highly variable and their primary sequences are substantially diverged). SC central region proteins from diverged organisms cannot necessarily be recognized as true homologs, but typically contain predicted coiled-coil domains and can collectively bridge the distance between the homolog axes (19).

CO recombination events are completed in the context of assembled SCs, yet we are only beginning to understand the complex interrelationships between the SC and COs. One conserved function of SC central region proteins is to promote normal levels of COs. However, organisms vary widely in the degree to which meiotic CO formation depends on SC central region proteins (20–23). Further, the pro-CO function(s) of these proteins may be exerted locally at CO sites, not necessarily requiring assembly of extended stretches of SC (24, 25). Conversely, SC central region proteins have also been implicated in antagonizing the formation of excess COs. This was first demonstrated in *C. elegans*, where essentially all COs are dependent on its SC central region proteins (SYP-1-4) and COs are normally limited to one per homolog pair (22, 26–28); whereas lack of SYP proteins eliminates interhomolog COs, altering SC composition by partial depletion of SYP proteins increases the number of COs and attenuates CO interference (29, 30). Recent work suggests that the SC central region may also play a role in limiting CO formation in *S. cerevisiae*, as CO numbers are similarly increased in mutants where the transverse filament protein Zip1p is present at recombination sites but cannot localize along the full length of a bivalent due to either an altered N-terminus or lack of other central region components (24). The observation that SC central region proteins can both promote and limit COs suggests the possibility that formation of CO intermediates would lead to a change in some assayable propert(ies) of the SC. This proposition would be difficult to test in experimental systems such as mouse or yeast, where assembly of the SC is coupled to and dependent on the formation of recombination intermediates. However, assembly of full length SCs between paired homologs in *C. elegans* is not dependent on recombination (26, 31), making this an ideal experimental system for investigating how formation of CO recombination intermediates might alter the state of the SC.

In this work, we investigate how SCs change during meiotic prophase progression in *C. elegans*. Our findings contribute to a growing body of evidence that the SC is a much more dynamic structure than is suggested by its highly-ordered appearance in EM images. We show that the SC incorporates new subunits and switches from a more dynamic to a more stable state during progression from the early pachytene to late pachytene stage. Moreover, we demonstrate that meiotic recombination events can indeed alter the state of the SC in a chromosome-autonomous manner.

## Results

### The SC changes during progression through the pachytene stage

The pachytene stage of meiotic prophase is defined by the presence of synaptonemal complex along the full interface between lengthwise aligned pairs of homologous chromosomes. However, composite fluorescence images of *C. elegans* gonads generated using an automated stitching algorithm ((32) which preserves relative fluorescence intensities of different fluorophores across a set of tiled images) reveal that nuclei that are in the pachytene stage based on this definition change in their appearance during the time that they spend in this stage. In immunofluorescence (IF) images of whole-mount gonads (Fig 1A), the IF signal intensity for SC central region protein SYP-1 gradually increases while the signal for chromosome axis protein HTP-3 remains relatively constant as nuclei progress through the pachytene stage. Further, in live worms expressing an EmGFP-tagged version of SC central region protein SYP-3 (Fig 1B), the EmGFP::SYP-3 fluorescence intensity in germ cell nuclei increases approximately twofold (2.1 ± 0.3, 2 gonads analyzed) over the course of pachytene progression. A similar increase in intensity also occurs in mutants lacking meiotic recombination (2.2 ± 0.1; Fig S1, 3 gonads analyzed) and presumably reflects the ongoing addition of SC central region subunits to existing “full length” SCs. Together, these observations indicate that the relative subunit composition and/or structure of SCs are not constant throughout the pachytene stage, but instead vary during pachytene progression.

**Figure 1.**
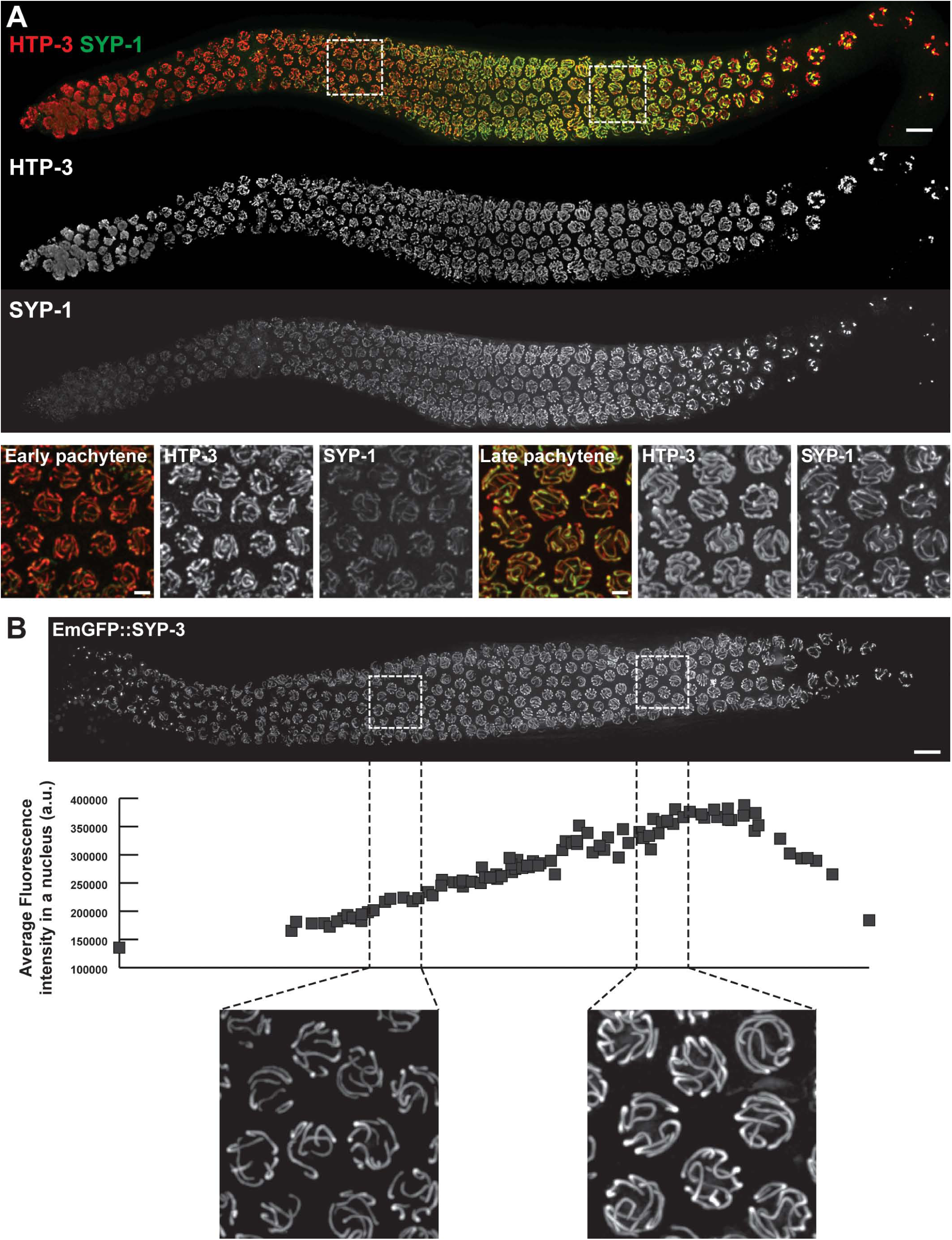
Levels of SC central region components SYP-1 and SYP-3 increase duringpachytene progression. **A.** Immuno-detection of wild-type distribution of SC components in a fixed whole-mount gonad (top) and higher magnification images of representative fields of nuclei in early (bottom left) and late pachytene (bottom right) stages. Chromosome axis protein HTP-3 is shown in red; SC central region protein SYP-1 is shown in green. **B.** Detection of SC central region component SYP-3 fused to GFP (EmGFP::SYP-3) in the gonad of a live worm (top). The graph (middle) represents the evolution of the average GFP::SYP-3 fluorescence intensity in a nucleus as a function of its position in the gonad (see Materials and Methods for details). The increase in GFP::SYP-3 signal during pachytene progression is illustrated in higher magnification fields of early and late pachytene nuclei (bottom). For both panels, images are maximum intensity projections of 3D data stacks encompassing whole nuclei. Scale bars represent 10 μm (whole gonad) and 2 μm (fields of nuclei).

Paralleling our observations in live worms and whole-mount dissected gonads, we also find that early and late pachytene nuclei differ in the sensitivity of their SCs to experimental perturbation by detergent-based lysis and partial spreading procedures (Fig S2–Fig S4). These observations reinforce the conclusion that the SC changes during progression through the pachytene stage of meiotic prophase.

### The SC central region switches from a more dynamic to a less dynamic state during pachytene progression

We investigated the potential dynamic nature of the SC central region using Fluorescence Recovery After Photobleaching (FRAP) in live worms expressing GFP-tagged versions of SC central region component SYP-3 (Fig 2). We used two distinct experimental systems and two independently-generated GFP::SYP-3 transgenic lines. For both transgenes, experiments were conducted using strains that also expressed untagged SYP-3 from the endogenous locus, as neither transgene fully rescued a *syp-3(null)* mutant (see Materials and Methods). Both of these approaches led to the same conclusions, namely that the SC is indeed a dynamic structure, and that the SC central region changes from a more dynamic to a less dynamic state during meiotic progression.

**Figure 2:**
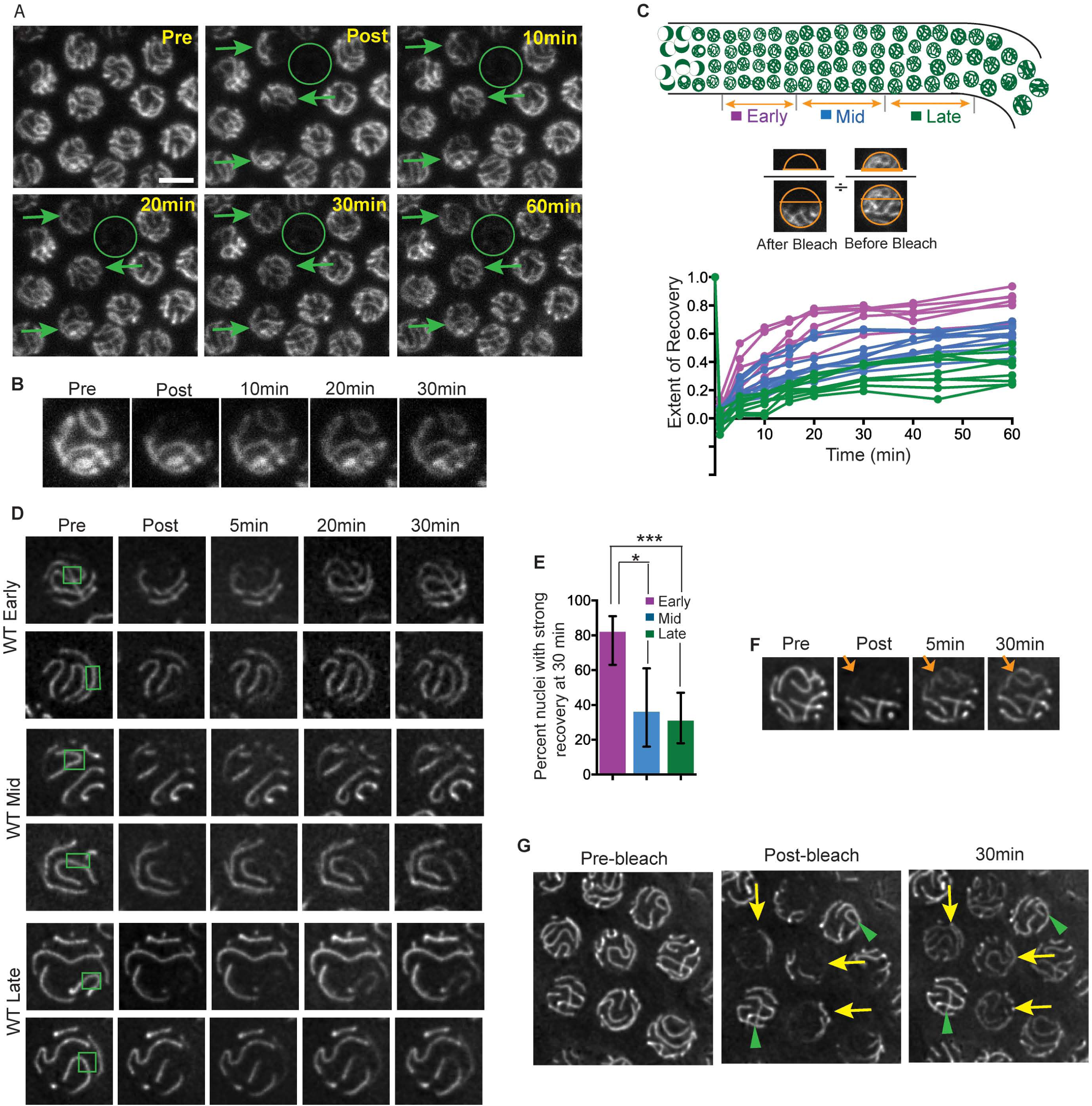
The SC switches from a more dynamic to a less dynamic state during pachytene progression. **A.** Confocal images of GFP::SYP-3 fluorescence from a FRAP experiment assessing fluorescence recovery in nuclei from the mid-pachytene region of a wild-type worm, following a Z-stack bleach protocol conducted using a Leica SP2 confocal microscope. Arrows indicate partially bleached nuclei and circle indicates a fully bleached nucleus; images are sum projections of image stacks encompassing whole nuclei. Scale bar is 5µm. **B.** Panels showing partial projection images depicting the upper half of a nucleus from A, illustrating clear recovery of fluorescence for an SC that was fully contained within the bleached portion of the nucleus. **C.** Quantitation of WT FRAP experiments performed using the confocal microscope (as depicted in A). Each curve in the graph plots the recovery of a single partially bleached nucleus, with the color indicating the zone (early, mid or late pachytene) of the scored nucleus, as depicted in the cartoon gonad scheme above the graph. Extent of recovery represents the ratio of the fluorescence (at a given post-bleach time point) in an ROI from the bleached portion of the nucleus to the pre-bleach fluorescence in that ROI, normalized to the total fluorescence in that nucleus. **D.** Panels of images of EmGFP::SYP-3 fluorescence in representative early, mid and late pachytene nuclei from FRAP experiments performed using the Deltavision OMX wide-field microscope. Photobleaching was focused at a single plane (in contrast to a stepwise Z-stack bleach), using small ROIs (1.2 - 3µm^2^) to bleach limited SC segments; green rectangles represent bleached regions. Images are partial projections showing the half of the nucleus containing the bleached SC segments. **E**. Quantitation of fluorescence recovery in early, mid and late pachytene nuclei for experiments of the type depicted in D (where strong recovery is defined as apparent evening-out of fluorescence between bleached and unbleached SC segments; see Materials and Methods for scoring methodology). *p = 0.0049; ***p < 0.0001; two-tailed Fisher exact test. **F.** Panel of EmGFP::SYP-3 images showing FRAP of an SC that had been bleached along its entire length (arrow), indicating that recovery involves redistribution of subunits among different SCs within a nucleus. **G.** GFP::SYP-3 images of a field containing a mixture of unbleached reference nuclei (green arrowheads) and nuclei in which a substantial portion of the SCs were bleached (yellow arrows). Fluorescence of the unbleached SC segments diminishes relative to unbleached reference nuclei, reflecting redistribution of subunits among the SCs within each bleached nucleus.

The experiments depicted in Fig 2A-C were carried out using a Leica SP2 confocal microscope in which the FRAP module was configured to perform a Z-stack bleach. In these experiments, a region corresponding to roughly 30-50% of each selected nucleus was subjected to stepwise photobleaching throughout a Z-stack (0.24 μm step size) encompassing the entire depth of the nucleus, and images were acquired at multiple subsequent time points (Fig 2A). For each nucleus analyzed, the extent of recovery at post-bleach time points was quantified by calculating the ratio of fluorescence signal within a Region of Interest (ROI) confined to the bleached portion of the nucleus relative to the fluorescence signal for that whole nucleus, normalized to the same ratio observed prior to the bleach (Fig 2C, S5A; Materials and Methods). Analyzed nuclei were grouped into “early”, “mid” and “late” pachytene stages based on their relative positions along the distal/proximal axis of the gonad. Data plotted in Fig 2C show that substantial recovery of fluorescence occurred, predominantly over a period of 30 minutes, with the majority of recovery achieved during the first 15-20 minutes. Moreover, extent of recovery was strongly correlated with the relative position of the nucleus within the pachytene region of the germ line (R^2^ = 0.71; Fig S5A), with the median recovery level observed for late pachytene nuclei being 51% lower than that observed for early pachytene nuclei (p <0.0001; Mann Whitney test). These data indicate that the SC central region becomes less dynamic as nuclei progress through the pachytene stage.

For Fig 2D-G, experiments were carried out using a Deltavision OMX Blaze wide-field deconvolution microscopy system, in which photobleaching is focused at a single selected focal plane (rather than bleaching stepwise throughout a Z-stack; see Materials and Methods). The images presented are partial projections showing the half of the nucleus containing the bleached portions of the SCs. For the experiments depicted in Fig 2D-E, small ROIs (1-3 μm^2^) were deliberately selected in order to bleach limited segments of the individual SCs within the nuclei; this approach allowed assessment of recovery in a context where the majority of the initial fluorescence in a nucleus was retained and potential photo-damage was minimized. Quantitation of these experiments (using a scoring system devised to accommodate movement of nuclei and chromosomes; see Materials and Methods) demonstrated that recovery was strongest in early pachytene nuclei, then diminished as nuclei progressed to the late pachytene stage (Fig 2E). Thus, FRAP analyses conducted using two very different experimental set-ups yielded the same findings, together providing strong support for the conclusion that SC transitions from a more highly dynamic state to a less dynamic state during pachytene progression.

Two additional conclusions can be made regarding the nature of the recovery observed in our experiments. First, several lines of evidence indicate that recovery occurs predominantly by redistribution of existing GFP::SYP-3 protein within a nucleus. Recovery was minimal in fully bleached nuclei, indicating that import of new protein from the cytoplasm is insufficient to account for the recovery observed during the time frame of the experiments (Fig 2A). (This is consistent with the that fact that the total amount of new fluorescent protein per nucleus is estimated to increase by less than 5% during the duration of a FRAP experiment, based on the observation that fluorescence increases by only two-fold during the entire course of the pachytene stage; Fig 1B.) Further, in our confocal experiments (where 30-60% of the initial fluorescence was eliminated by bleaching) the normalized total fluorescence within such a partially-bleached nucleus (relative to a neighboring unbleached reference nucleus) remained almost constant during the recovery time course (Fig S5B). Finally, in experiments using large ROIs, the unbleached portions of SCs within partially-bleached nuclei diminished in fluorescence relative to unbleached neighboring nuclei in the same field (Fig 2G).

A second conclusion can be drawn from images of nuclei in which an individual SC was bleached throughout its entire length (*e.g.* Figs 2B and 2F, S5C). Such SCs can nevertheless exhibit substantial recovery of fluorescence. This observation indicates that while recovery likely involves redistribution of subunits within individual SCs, it also involves redistribution of subunits among the different SCs within a nucleus, presumably through exchange with a nucleoplasmic protein pool.

### The more dynamic state of the SC is prolonged in mutants that cannot form COs

The observed transition in the dynamic state of the SC occurs in parallel with the process of meiotic recombination. Further, previous studies have revealed that cytological features characteristic of early pachytene are prolonged in mutants that are impaired in meiotic recombination (33-35). Thus, we conducted FRAP experiment to evaluate the dynamic state of the SC in *spo-11* mutants, which are defective in forming the DSBs that serve as the initiating events of recombination (31), and in *cosa-1* mutants, which are defective in converting resected DSBs into interhomolog crossovers (COs) (12) (Fig 4). Both the wide-field small ROI FRAP assay (Figs 3A, Figs 3C) and the confocal Z-stack bleach FRAP assay (Fig 3B) yielded the same overall findings. First, the substantial decline in recovery observed during progression from early to late pachytene in the wild type was not observed in *spo-11* and *cosa-1* mutants (albeit a modest 9% decline was detected in the confocal assay for the *cosa-1* mutant). Second, the degree of fluorescence recovery in late pachytene nuclei in both the *spo-11* and *cosa-1* mutants was significantly higher than in wild type. Thus, an inability to form CO recombination intermediates is associated with persistence of a more dynamic state of the SC. This suggests that formation of CO recombination intermediates may serve as a trigger for the SC to switch to a less dynamic state.

**Figure 3:**
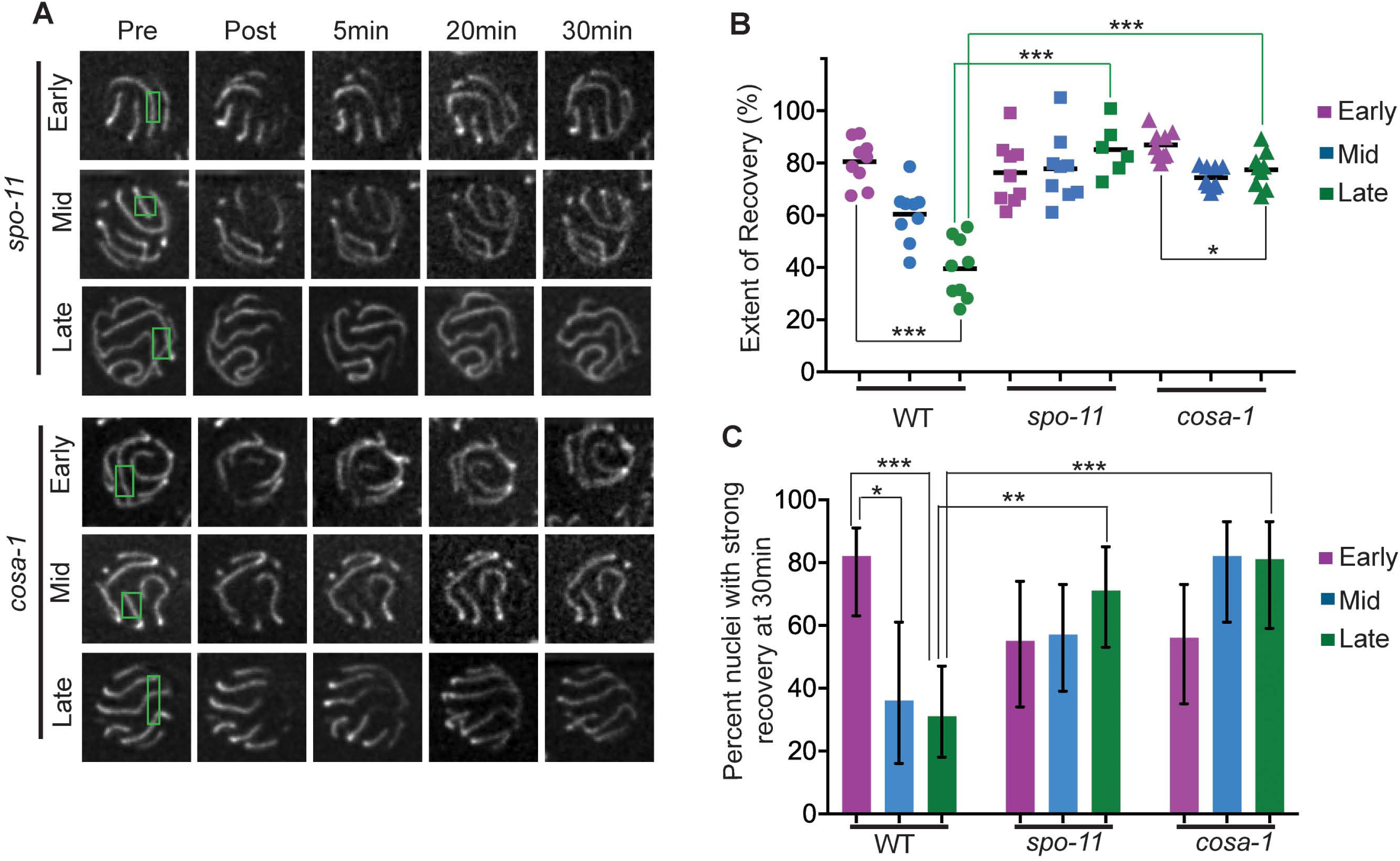
The more dynamic state of the SC is prolonged in mutants that cannot form COs. **A.** Panels of images of representative pachytene nuclei from the *spo-11* and *cosa-1* mutant, from FRAP experiments performed as in. Green rectangles indicate bleached regions. **B.** Quantitation of extent of fluorescence recovery in FRAP experiments conducted as in Fig 2A-C. Each data point represents the plateau value for the “extent of fluorescence recovery” curve for one nucleus. Whereas the extent of recovery in wild type was reduced by 51% between early and late pachytene in wild type, no significant decline in recovery was observed in the *spo-11* mutant and only a modest 9% decline was observed in the *cosa-1* mutant. Moreover, late pachytene nuclei in both of these mutants exhibited significantly higher recovery than in wild type. *p = 0.0056; ***p < 0.0005; two-tailed Mann-Whitney test. **C.** Quantitation of recovery in early, mid and late pachytene nuclei for FRAP experiments of the type depicted in A (as in Fig 2E) *0.01 < p < 0.05; **p = 0.0013; ***p < 0.0005; two-tailed Fisher exact test; for the *spo-11* and *cosa-1* mutants, no significant differences were observed between early, mid and late pachytene.

**Figure 4:**
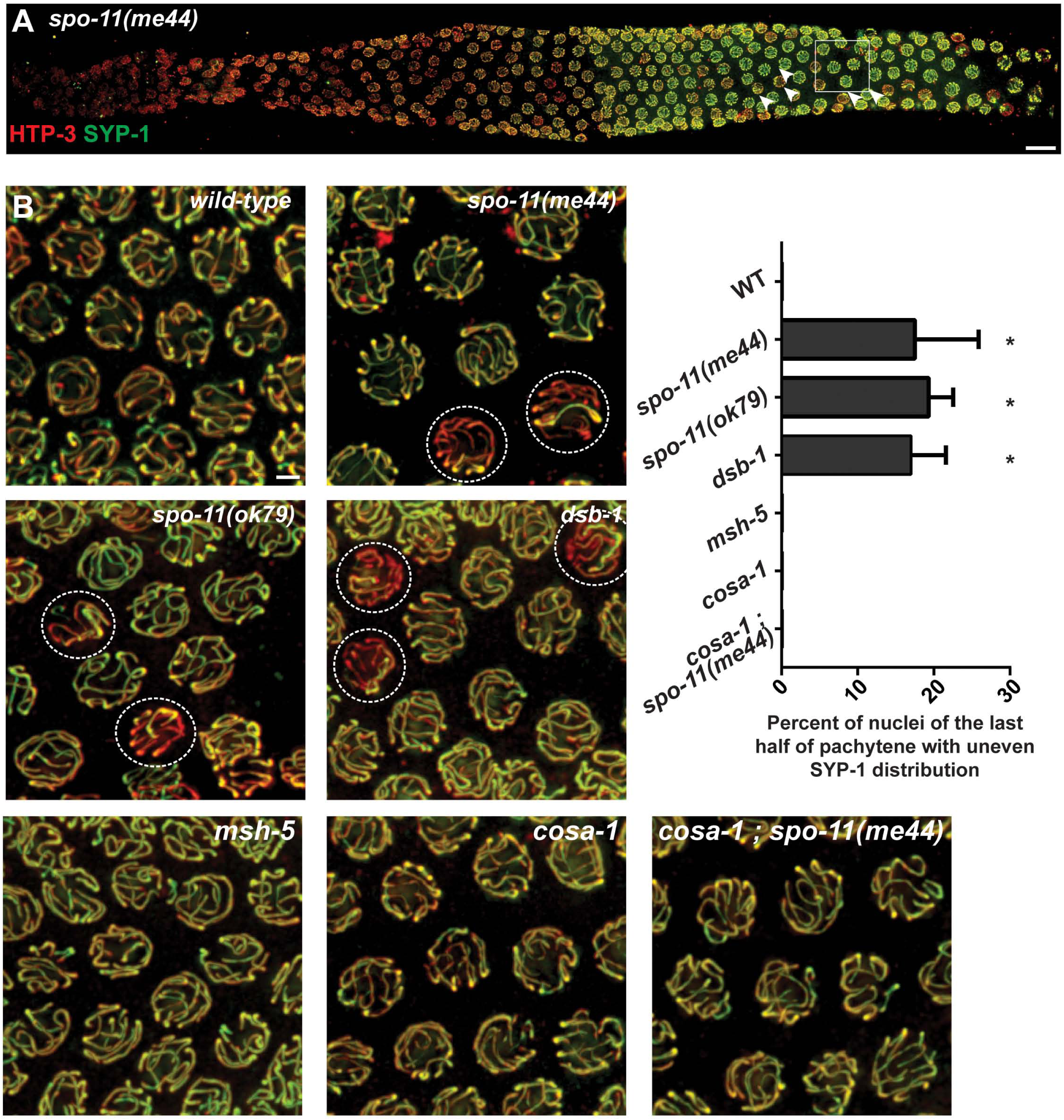
A subset of late pachytene nuclei in DSB-defective mutants exhibit an uneven distribution of SYP-1 among their SCs. **A.** Immunodetection of SC components HTP-3 (red)and SYP-1 (green) in a fixed whole-mount *spo-11(me44)* mutant gonad. In late pachytene, some nuclei, indicated by an arrowhead, display an uneven distribution of the central region component SYP-1. **B.** Representative fields of late pachytene nuclei observed in wild-type, in recombination initiation mutants *spo-11(me44)*, *spo-11(ok79)* and *dsb-1(tm5034)*, in the *msh-5(me23), cosa-1(tm3298)* mutants (which are proficient for initiation of recombination but defective in conversion of processed DSBs into COs), and in the *cosa-1(tm3298); spo-11(me44)* double mutant, stained for HTP-3 (red) and SYP-1 (green). Dashed circles indicate nuclei in the mutants with uneven SYP-1 distribution. For both panels A and B, images are maximum intensity projections of 3D data stacks encompassing whole nuclei. Scale bars represent 10μm (whole gonad, panel A) and 2μm (fields of nuclei, panel B). **C.** Quantification of the fraction of nuclei with uneven SYP-1 distribution in the last half of pachytene for each indicated genotype. Asterisks indicates highly significant differences (p < 0.0001; chi-square test) when single mutants are compared to wild-type or when *spo-11(me44)* is compared to *cosa-1(tm3298); spo-11(me44)*.

### A subset of nuclei in DSB-defective mutants exhibit an uneven distribution of SYP-1 among SCs

In the course of analyzing immunofluorescence images of whole-mount gonads from *spo-11* mutant worms, which lack the SPO-11 enzyme responsible for making meiotic DSBs, we noticed a difference compared to wild-type in the appearance of the SC immunostaining in the second half of the pachytene region (Fig 4): whereas SYP-1 immunostaining appeared uniformly distributed among the individual SCs in most wild-type nuclei, a subset of nuclei in *spo-11* mutant worms (17 ± 7% for *spo-11(me44)*, 20 ± 3% for *spo-11(ok79)*) exhibited an uneven distribution of SYP-1 among the SCs, and within the last quarter of the pachytene region, it was clear that SYP-1 staining was preferentially enriched on a single SC relative to the other SCs in that same nucleus. Late pachytene nuclei with uneven SYP-1 distribution were likewise observed in the *dsb-1* mutant (20 ± 3%), which is also impaired in initiation of meiotic recombination (34), indicating that the presence of such nuclei is a shared property of mutants that are proficient for synapsis but defective in the formation of SPO-11-dependent DSBs.

Both the incidence of late pachytene nuclei exhibiting uneven SYP-1 distribution (p<0.0001, Fisher’s exact test) and the number of SYP-1-enriched SCs within such nuclei (p<0.0001, Mann Whitney test) increased following low dose IR treatment of the *spo-11* mutant, suggesting that in this context, DNA breaks can trigger enrichment of SYP-1 proteins on a subset of chromosomes (Fig S6). Further, the number of SYP-1-enriched SCs in late pachytene nuclei was significantly lower (p<0.0001, Mann Whitney test) in IR-treated *spo-11* worms heterozygous for a reciprocal translocation (*spo-11; szT1(I;X)/+*), in which approximately 33% of the genome is engaged in heterosynapsis, than in IR-treated *spo-11* worms (Fig S6). This further suggests that the ability of IR-induced DNA breaks to trigger SYP enrichment depends on the ability of such breaks to engage in recombination-based interactions with homologous DNA sequences.

Our finding that IR-induced DNA breaks produced in limiting numbers can elicit uneven distribution of SYP-1 within nuclei seems at odds with the fact that nuclei with uneven SYP-1 distribution occur even in the absence of irradiation in the meiotic-DSB-defective mutants. These results can be reconciled by hypothesizing that spontaneous SPO-11-independent DNA lesions (of unknown structure) occur at a low frequency in mutants lacking meiotic DSBs. While such spontaneous DNA lesions do not appear to yield inter-homolog COs (31), they are nevertheless capable of recruiting meiotic DNA repair proteins when the normal preferred substrates for these proteins are absent (see below).

### Chromosomal enrichment of SYP-1 reflects the recruitment of pro-CO factors COSA-1 / MutS_γ_

Two lines of evidence support the conclusion that the uneven distribution of SYP-1 protein within nuclei in the context of limiting DNA breaks reflects the formation of recombination/repair intermediates that recruit meiotic CO factors.

First, the presence of nuclei exhibiting an uneven distribution of SYP-1 requires the presence pro-CO factors COSA-1 and MutS_γ_ (MSH-4–MSH-5) (Fig 4), as such nuclei are not detected in *cosa-1* or *msh-5* single mutants, which are proficient for DSB formation but defective in repairing DSBs as COs. Further, such nuclei are not detected in *spo-11; cosa-1* double mutants, indicating that COSA-1 is required to trigger the uneven SYP-1 distribution observed in a *spo-11* mutant background.

Second, when DSBs are limiting, SYP-1 is preferentially concentrated on the subset of SCs harboring a focus where pro-CO factors are concentrated (Fig 5A-D). Late pachytene nuclei in wild-type worms contain 6 bright GFP::COSA-1 foci marking the site of the single CO on each homolog pair (12). Most late pachytene nuclei in *spo-11* worms expressing GFP::COSA-1 completely lack COSA-1 foci, consistent with an absence of DSBs and CO recombination intermediates (12). However, a subset of late pachytene nuclei in these worms harbor a single COSA-1 focus, suggesting the infrequent occurrence of a SPO-11-independent DNA lesion that is capable of recruiting meiotic recombination factors during the late pachytene stage. Further, whereas late pachytene nuclei lacking COSA-1 foci exhibited a uniform distribution of SYP-1, nuclei with a COSA-1 focus exhibited uneven SYP-1 distribution, with SYP-1 becoming preferentially enriched on the SC harboring the COSA-1 focus (Fig 5A).

**Figure 5:**
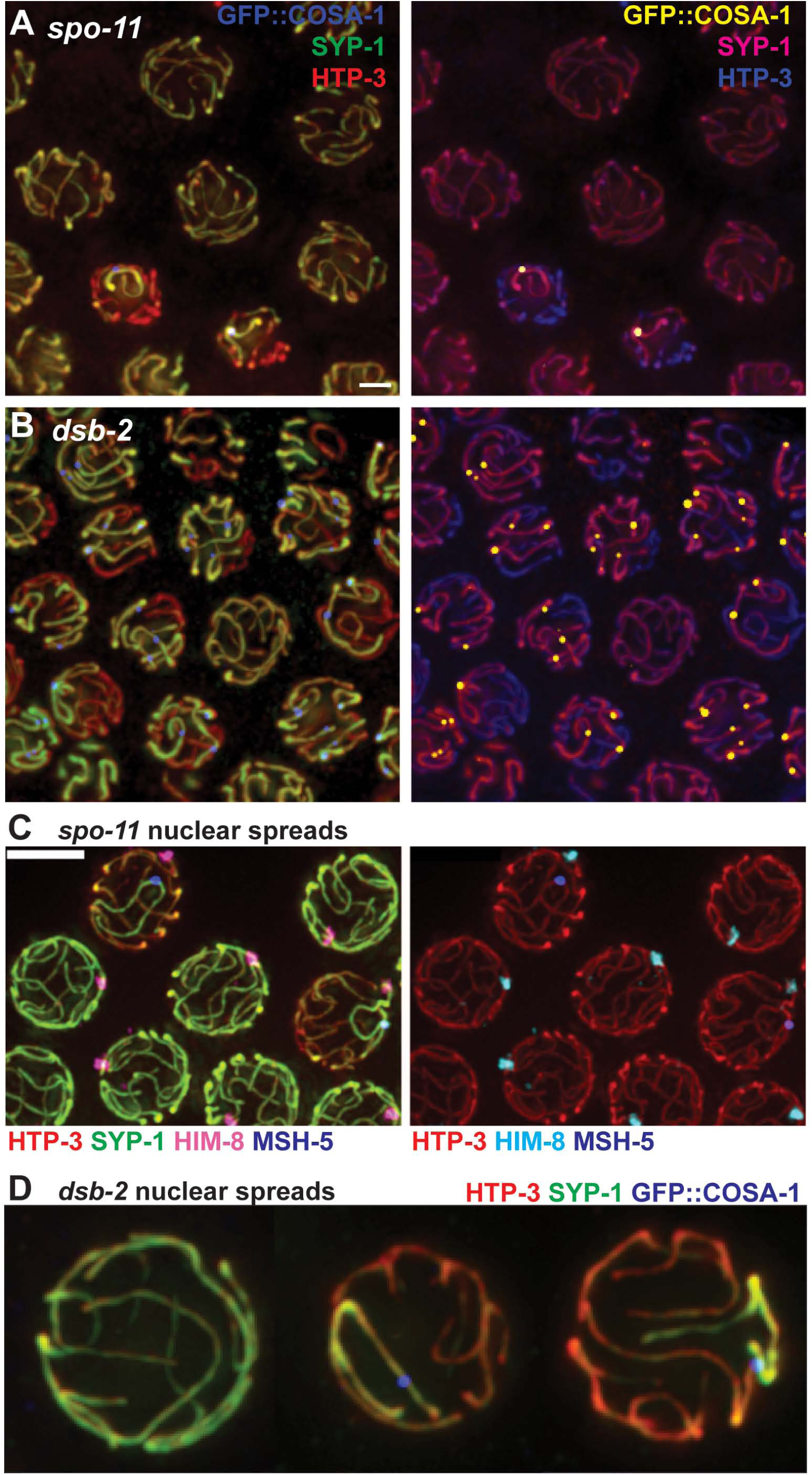
When DSBs are limiting, SYP-1 is preferentially enriched on bivalents with a site where pro-crossover factors COSA-1 and MutS_γ_ are concentrated. **A and B.** Representative fields of whole-mount late pachytene nuclei in recombination initiation mutants *spo-11(me44)* and *dsb-2(me96)* immunostained for HTP-3, SYP-1 and GFP::COSA-1. Images are shown in two different color schemes to emphasize the correspondence between the presence of a GFP::COSA-1 focus and enrichment of SYP-1 on a chromosome pair. Scale bar represents 2μm. **C.** Representative field of spread nuclei at the early-to-late pachytene transition from the gonad of a *spo-11(me44)* mutant worm grown at 25°C and immunostained for HTP-3, SYP-1, MSH-5 and HIM-8. HIM-8 was used identify the X chromosomes, as it marks the pairing center locus located near the left end of X. In the nuclei containing a MSH-5 focus, SYP-1 is enriched on the chromosome pair harboring the focus. The SYP-1-enriched chromosome with a recombination focus can either be the X chromosome (middle right) or an autosome (upper left). In the remaining nuclei, no chromosome displays a recombination focus and SYP-1 is distributed equally among all chromosomes. (For this experiment, worms were raised at 20°C until the late L4 stage, then shifted to 25°C for 24 h prior to dissection; under these conditions, the incidence of nuclei with a SYP-1-enriched chromosome harboring a recombination focus is higher than at 20°C.) Scale bar is 5μm. D. Isolated late pachytene nuclear spreads from the *dsb-2(me96)* mutant, illustrating even distribution of SYP-1 among all chromosomes in anucleus lacking a GFP::COSA-1 focus (left) and enrichment of SYP-1 on the single chromosome harboring a COSA-1 focus in the middle and right nuclei.

Experiments using MSH-5 as a recombination marker gave identical results. (Fig 5C). These experiments further showed that when a single SYP-1-enriched SC harboring a recombination focus is detected, it can be associated either with the X chromosomes or with an autosome pair (Fig 5C). However, we note that association with the X chromosomes is disproportionate (34% observed vs. 17% expected; n = 38, p = 0.0018), perhaps reflecting known differences in replication timing (36) and/or chromatin structure (37) between the X and autosomes that might affect the incidence of the triggering DNA lesions.

Analysis of late pachytene nuclei in a *dsb-2* mutant, in which SPO-11-dependent DSBs are substantially reduced but not eliminated (33), strongly corroborated the correspondence between the presence of COSA-1 foci in late pachytene nuclei and uneven distribution of SYP proteins (Fig 5B, D). Likewise, within such nuclei, SYP-1 was enriched on the subset of SCs harboring the COSA-1 foci.

Taken together, our data indicate that when DSBs are limiting, concentration of CO proteins at a putative DNA repair site triggers a change in status of SC components, resulting in enrichment of SYP proteins in *cis* on the chromosome harboring such an event. We conclude that COSA-1-marked recombination intermediates, and/or concentration of pro-CO factors at such sites, cause chromosome-autonomous stabilization of the SCs.

### Polo-like kinase PLK-2 as a candidate mediator of recombination-triggered change in SC state

To better understand how formation of a COSA-1 / MutS_γ_ dependent recombination intermediate (and/or recruitment of these pro-CO factors) triggers a change in state of the SC with which it is associated, we investigated the potential involvement of polo-like kinase PLK-2. Whereas PLK-2 is initially concentrated at pairing centers (nuclear envelope-associated chromosome sites that coordinate chromosome movement, homolog recognition and SC assembly) during early prophase, it exhibits dynamic localization during meiosis and is subsequently detected on the SCs later during the pachytene stage (38, 39).

We first re-examined PLK-2 localization during meiotic prophase progression in germ cell nuclei from otherwise wild-type worms expressing PLK-2::HA from the endogenous *plk-2* locus, prepared using mild detergent-based lysis conditions (Fig 6, top). Our images refine the previously reported dynamic localization pattern of PLK-2, showing in addition that: 1) localization of PLK-2 on SCs can be detected very early in pachytene, and 2) preferential concentration of PLK-2 on limited SC subdomains (defined by recombination sites) can be detected right after the early-to-late pachytene transition, substantially earlier than the SYP proteins exhibit a similar pattern of CO-triggered relocalization. This suggests that PLK-2 localization during pachytene progression may not simply track with the SYP proteins, but may predict the behavior of SYP proteins during late pachytene.

**Figure 6:**
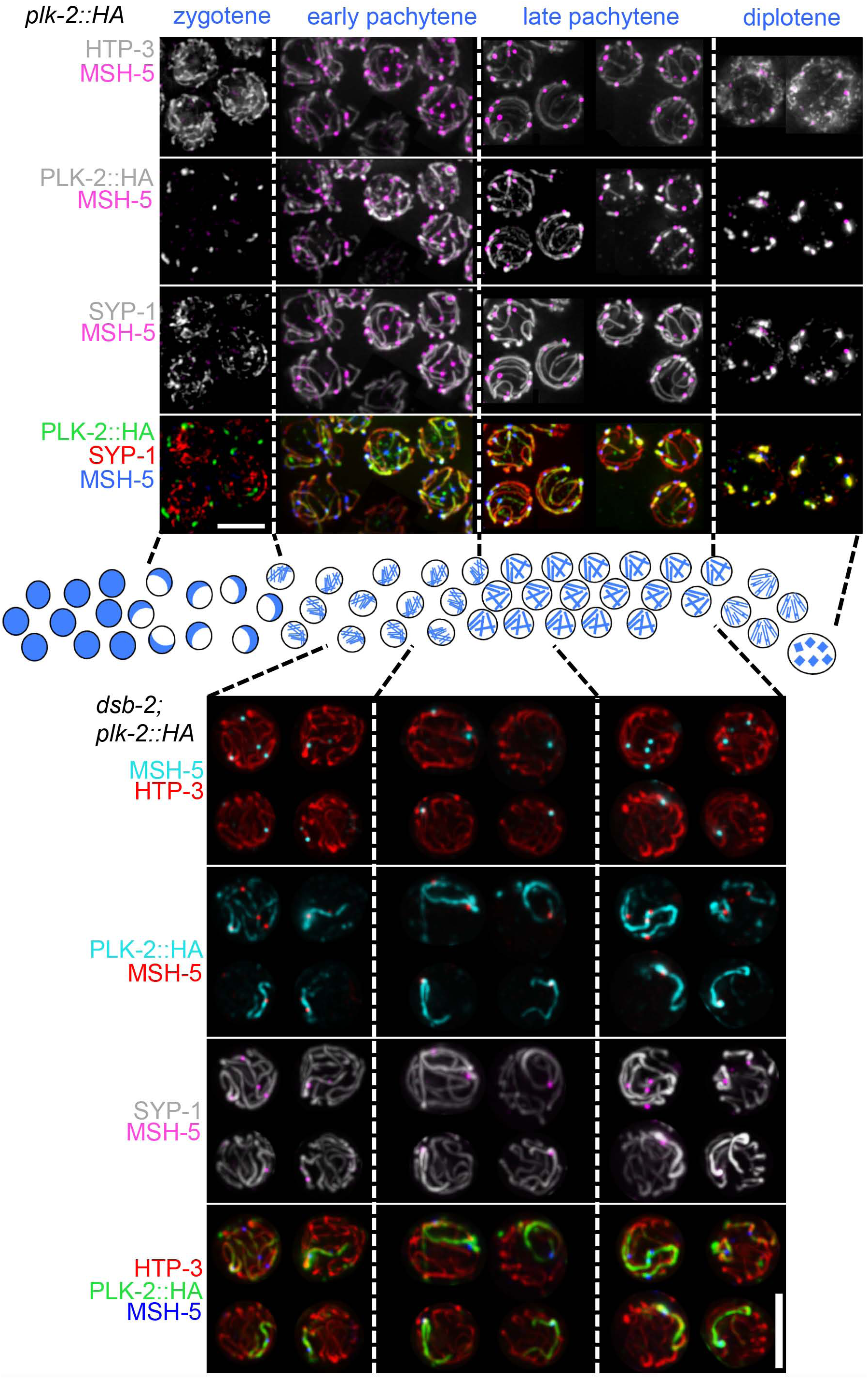
Localization of PLK-2 with meiotic chromosomes is highly dynamic and is influenced by recombination. **Top:** Dynamic localization of PLK-2 during wild-type meiosis.Representative images of spread nuclei from each indicated stage (zygotene, early pachytene, late pachytene, diplotene) in wild-type worms expressing HA-tagged PLK-2. As previously reported (Harper et al., 2011 and Labella et al., 2011), PLK-2 localizes to the pairing centers at the nuclear envelope during zygotene, while homolog pairing is being established and SCs are assembling, In early pachytene, PLK-2 becomes localized along synapsed bivalents that are engaged in recombination (*i.e.* displaying at least one MSH-5 focus). Prior to the early-to-late pachytene transition, PLK-2 localizes along the whole length of a bivalent. After the early-to-late pachytene transition, PLK-2 starts to retract, eventually becoming restricted to one side of the nascent CO site (marked by MSH-5); this restriction of PLK-2 to the “short arm” of the bivalent can be detected prior to detection of a similar retraction of SYP-1 that becomes visible by the end of pachytene (see late pachytene panels). By diplotene, both PLK-2 and SYP-1 are concentrated around chiasmata and on the short arm. **Bottom:** Recombination-triggered relocalization of PLK-2 is detected prior to SYP-1 enrichment on chromosomes undergoing recombination. Representative images of spread nuclei from each indicated stage (early pachytene, mid pachytene and late pachytene) from a *dsb-2* mutant expressing PLK-2::HA. During early pachytene, PLK-2 is first observed to be enriched on SCs near recombination sites marked by a MSH-5 focus and then becomes concentrated along the whole length of the recombining bivalent. Full-length localization of PLK-2 along a bivalent with a MSH-5 focus is detected prior to similar enrichment of SYP-1, which is first detected in mid pachytene. In nuclei lacking recombination events, PLK-2 is distributed uniformly among the bivalents (not depicted), as observed for SYP-1. Note: PLK-2 localizes not only to nuclei, but also to other extra-nuclear structures in the gonad, such as the rachis walls; consequently, IF signals from these other structures will tend to obscure the nuclear PLK-2 signals in maximum intensity projections of fields of nuclei. Thus, to facilitate visualization of subnuclear localization of PLK-2, representative spread nuclei were individually cropped from the edges of whole gonad spreads. In addition, the MSH-5 signal was overlaid as a screen on top of the other channels in order to facilitate visualization of the locations of MSH-5 foci in the projected images. Scale bar is 5μm.

Analysis of PLK-2 localization in the *dsb-2* mutant clearly shows that PLK-2 becomes highly enriched on the subset of chromosomes harboring a MutS_γ_-marked recombination site (Fig 6, bottom). Moreover, strong preferential localization of PLK-2 on MutS_γ_-marked chromosomes is clearly detected much earlier than preferential concentration of SYP-1 is detected on such chromosomes. These findings are consistent with a model in which COSA-1/ MutS_γ_- marked recombination sites trigger relocalization of PLK-2, which in turn affects the state of the SC.

To complement these localization experiments showing that PLK-2 is present at the right time and place to mediate changes in SC state, we conducted experiments using the null mutation *plk-2(ok1336)* to test the impact of loss of PLK-2 function on the state of the SC during the late pachytene stage (Fig 7). The early meiotic roles of PLK-2 in promoting homolog pairing and synapsis are substantially supplanted in the *plk-2(ok1336)* mutant by its close paralog PLK-1; thus, while SC assembly is delayed in this mutant, most chromosome regions are synapsed by late pachytene and most chromosome pairs recruit COSA-1 foci and form chiasmata (38, 39) (Fig 7C).

**Figure 7:**
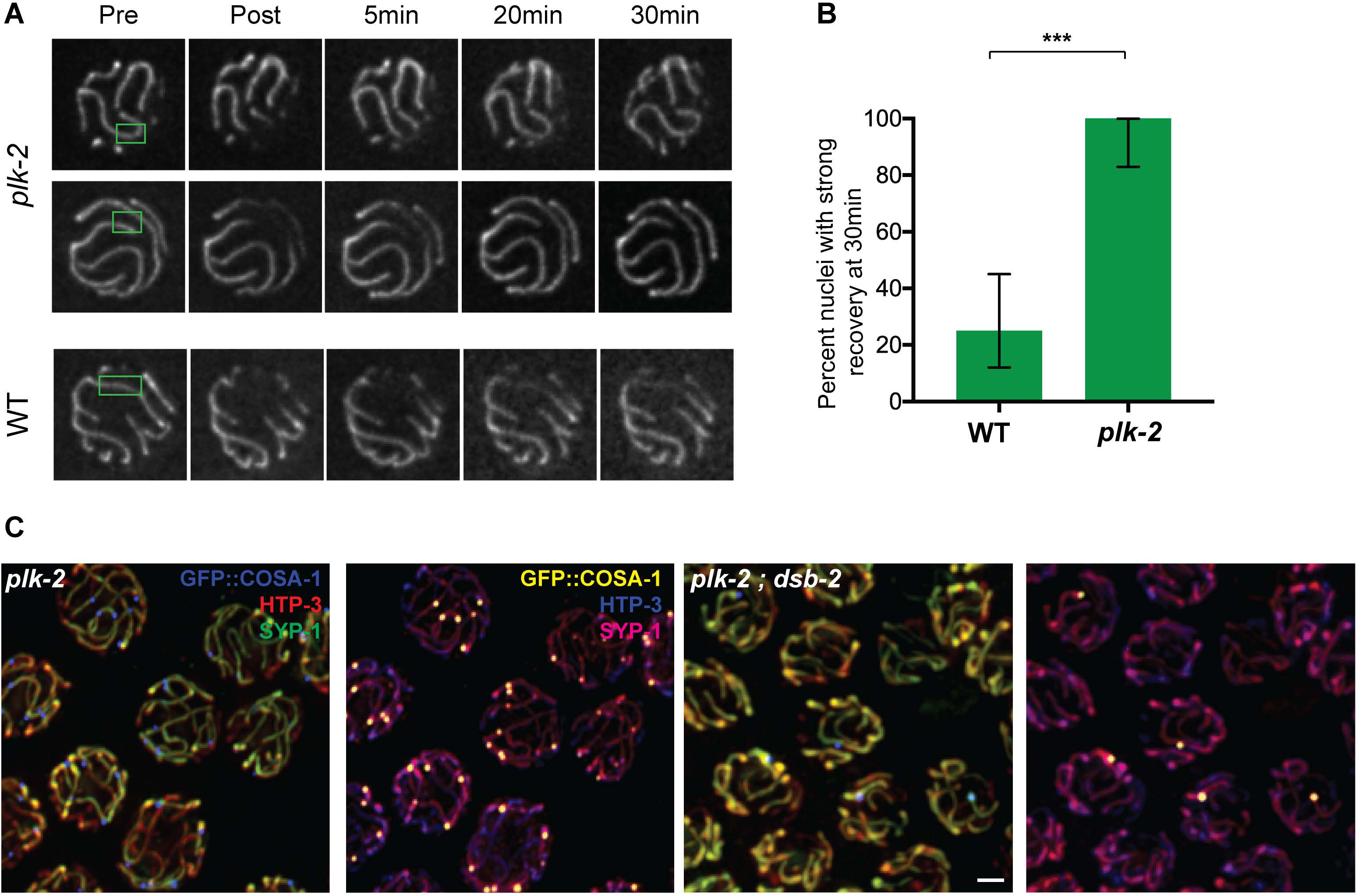
PLK-2 is required both for normal late pachytene SC dynamics and for preferential enrichment of SYP-1 on COSA-1-marked chromosomes when DSBs are limiting. **A.** Panels of images of EmGFP::SYP-3 fluorescence in representative late pachytene nuclei from FRAP experiments performed as in Fig 2B, illustrating the strong fluorescence recovery that was consistently observed in the *plk-2* mutant. **B.** Quantitation of fluorescence recovery in late pachytene nuclei from wild-type and *plk-2* mutant worms (as in Fig 2E, Fig 3C). ***p < 0.0001; two-tailed Fisher exact test. **C.** Immunofluorescence images of late pachytene nuclei from the *plk-2(ok1336)* single mutant and *plk-2(ok1336); dsb-2(me96)* double mutant; images are maximum intensity projections of 3D data stacks encompassing whole nuclei treated as in Fig 5A–Fig 5B. Scale bar represents 2μm. In contrast to the *dsb-2* single mutant (Fig 5B), SYP-1 signals are not specifically enriched on the chromosomes with a COSA-1 focus in the *plk-2;dsb-2* double mutant. (Note that there are asynapsed chromosome segments present in somenuclei, reflecting partially impaired synapsis in the *plk-2* mutant background.)

Fig 7A-B show the results of FRAP experiments comparing GFP::SYP-3 dynamics in late pachytene nuclei of wild-type and *plk-2* mutant worms. Strong fluorescence recovery was consistently detected in all late pachytene nuclei in the *plk-2* mutant background, reflecting a more highly dynamic state of the SC central region in the *plk-2* mutant than in wild-type controls. Similarly, immunofluorescence analysis of late pachytene nuclei in the *plk-2; dsb-2* double mutant revealed that in contrast to the *dsb-2* single mutant (Fig 5), SYP-1 does not become preferentially concentrated on the subset of chromosomes that harbor a COSA-1-marked recombination site (Fig 7C).

Together, our data show that 1) PLK-2 becomes localized on chromosomes harboring MutS_γ_-marked recombination intermediates in a manner that precedes and predicts the later localization of SYP proteins, and 2) PLK-2 is required to elicit changes in state of the SC that are normally triggered by such intermediates.

## Discussion

Accumulating evidence from experiments where the meiotic program had been experimentally challenged has contributed to a growing recognition the SC is a more dynamic structure than is suggested by its highly-ordered appearance in EM images. The first hint came from analysis of meiosis in mouse spermatocytes with homologs of different lengths (because of heterozygosity for a chromosome rearrangement), where regions of asynapsis were observed in early pachynema but not detected in late pachynema, suggesting that a process of “synaptic adjustment” operates to minimize asynapsis (40). Synaptic adjustment has likewise been demonstrated to occur during *C. elegans* meiosis (41, 42). More recently, investigation of pachytene arrested cells in *S. cerevisiae* demonstrated that SC central region subunits are continually incorporated into previously assembled SCs (43). Further, analysis of synaptic configurations in *C. elegans* polyploid meiocytes revealed that synaptic connections between more than two homologs were detected in early pachynema, whereas late synaptic interactions were invariably pairwise (44), suggesting that the properties of the SC change upon pachytene progression.

In the current work, we clearly demonstrate dynamic features of the *C. elegans* SC is that can be observed not only when the meiotic program is experimentally perturbed, but also in the context of normal meiosis. Over the normal course of progression through the pachytene stage, SCs undergo ongoing incorporation of additional central region (SYP) subunits, become increasingly resistant to disruption by detergent lysis/spreading procedures, and transition from a more dynamic to a less dynamic state as assessed by FRAP analysis. Further whereas the ongoing incorporation of SYP subunits is not dependent on recombination, we demonstrate that recombination is important for the stabilization of SC structure revealed by FRAP analysis.

### Recombination-triggered modulation of SC state

We have presented two independent lines of evidence that converge on the conclusion that formation of CO-eligible recombination intermediates triggers a change in state of the SC. First, our FRAP analyses revealed that a more highly dynamic status of the SC central region characteristic of early pachynema in wild-type meiosis is retained in late pachytene nuclei both in the *spo-11* mutant, which fails to initiate recombination (31), and in the *cosa-1* mutant, which fails to accumulate pro-CO factors at recombination sites and can’t form COs (12). These data indicate that proper execution of meiotic recombination is responsible for at least part of the change in dynamics observed during prophase progression.

Second, whereas FRAP analysis indicated a link between recombination and SC stabilization, IF analyses of SCs under conditions where meiotic DSBs are limiting helped to clarify this relationship. These IF experiments revealed that when chromosomes with and without a COSA-1-marked (or MutS_γ_ -marked) recombination intermediate co-exist within the same nucleus, SYP proteins become preferentially associated with the chromosome pair(s) that harbor such a recombination intermediate, apparently at the expense of non-recombinant chromosomes. In light of our observations in the FRAP analyses, this latter set of findings supports the view that the effects of recombination on SYP dynamics are not nucleus-wide but are instead exerted at the level of individual chromosome pairs; *i.e.,* we infer that CO recombination events alter the SC central region in chromosome-autonomous manner.

Together our data allow us to conclude that SC central region stabilization is triggered either by a recombination intermediate that is created and/or stabilized by MSH-5 and COSA-1, or by the physical presence of these pro-CO proteins at recombination sites. We further reasoned that homologous engagement is necessary, but not sufficient, for these changes in the properties of the SC: The need for homolog engagement can be inferred from our experiments showing that the incidence of SYP-enriched chromosomes detected following induction of DNA breaks by IR in *spo-11* mutant meiocytes is significantly reduced when one-third of the genome is engaged in heterologous (rather than homologous) synapsis. Conversely, the insufficiency of homolog engagement is implied by the fact that pro-CO factors are required for SC central region stabilization despite the fact that *msh-5* mutants are competent to engage in homologous recombination to repair DSBs induced during meiosis as inter-homolog non-COs (45).

### Coupling CO intermediate formation to SC stabilization

How can a nascent CO event at one site on a chromosome pair trigger a chromosome-wide change in the properties of the SC? Our experiments implicate the polo-like kinase PLK-2 as a key player in this process. Polo-like kinases are known to play multiple roles during mitosis and meiosis. For example, the POLO homolog Cdc5 is required both to resolve meiotic CO intermediates and to disassemble the SC at pachytene exit during budding yeast meiosis (46). In *C. elegans*, previous work has implicated PLK-2 in multiple meiotic functions, including: 1) promoting early prophase chromosome movements coupled to homolog pairing and SC assembly, 2) checkpoint responses to unsynapsed chromosomes, and 3) normal restructuring of bivalents during late meiotic prophase (38, 39). Based on our current data, we propose a new role, in coupling formation of CO-eligible recombination intermediates to stabilization of the SC central region.

Our thinking is shaped by integration of findings from functional assays and from PLK-2 protein localization. FRAP analysis showed that late pachytene nuclei in a *plk-2* mutant exhibit a more highly dynamic SC state than in wild type germ lines, and IF analysis showed that PLK-2 is required to achieve preferential association of SYP-1 on chromosomes harboring CO sites (when recombination intermediates are present in limiting numbers). On their own, these findings are consistent either with a) PLK-2 being required for the SC central region to mature to a state where CO intermediates are capable of triggering a change in SC state, or with b) PLK-2 playing a role in eliciting the state change. Our analysis of PLK-2 localization under conditions where DSBs (and consequent CO intermediates) are limiting leads us to favor the latter interpretation. Concentration of PLK-2 along the length of chromosomes harboring CO sites is detected prior to the eventual enrichment of SYP proteins on such chromosomes. This observation, together with the requirement for PLK-2 to achieve this preferential SYP association, are best explained by a model in which nascent CO sites serve to recruit PLK-2 to the SCs that harbor them. We hypothesize that PLK-2 localization and activity then spread along the lengths of these SCs via a self-reinforcing feedback loop that in turn results in stabilization of the SC central region along the full length of the SC. This model is reinforced by the recent finding of S. Nadarajan and M. Colaiacovo (personal communication) that SC central region protein SYP-4 is phosphorylated in a PLK-1/2-dependent manner.

### Roles of CO-intermediate-triggered SC stabilization

What function(s) might be served by coupling formation of CO-eligible recombination intermediates to a change in state of the SC? We can envision several possible mechanisms by which such a coupling could help to promote a robust outcome of meiosis:

First, we propose that CO-intermediate-triggered alteration of SCs may play a key role in a regulatory/surveillance network that regulates meiotic progression and promotes CO assurance (33–35). This surveillance network monitors the formation of CO-eligible recombination intermediates and makes timely progression from early pachytene to late pachytene contingent on their occurrence. Progression to late pachytene is characterized by a major coordinated transition affecting multiple distinct aspects of the meiotic program, including cessation of DSB formation, maturation of CO sites, and loss of the ability to engage the homolog as a template for DNA repair.

A key feature of this surveillance network is that germ cells can detect a single chromosome pair lacking a CO intermediate, prolonging the early pachytene stage in response. However, it was difficult to envision how a chromosome pair lacking a relevant recombination intermediate might be detected. Based on our current findings, we propose that the state of the SC may in fact be the monitored feature, and that CO-triggered modulation of SC state could serve as a means to extinguish a signal that maintains the early pachytene stage. According to this model, formation of the required CO-eligible intermediates on all chromosome pairs would lead to all six SCs becoming stabilized, which would in turn extinguish the “maintain early pachytene” signal and thereby enable nuclei to progress to the late pachytene stage.

The model proposed here is distinct in several ways from the model recently proposed by Machinova and colleagues (47). Machinova *et al.* similarly found that under conditions where DSBs are limiting, SC central region proteins are preferentially retained or stabilized on chromosomes undergoing recombination events. Further, using genetic analyses complementary to those reported here, they likewise implicated COSA-1, MutS_γ_ and PLK-2 in this process. In their model, Machinova *et al.* asserted that nuclei respond to detection of synapsed chromosome pairs that lack CO intermediates by actively triggering desynapsis/destabilization of the SC in late pachytene to provide another chance to form COs. Given our demonstration that the SC central region exhibits a more highly dynamic state in early pachytene that is shut down during progression to late pachytene, we argue that it is not necessary to invoke active desynapsis to explain the data. Rather, we suggest that the reduction of SYP proteins on late pachytene bivalents lacking CO sites may be a secondary, pathological, consequence of SYPs becoming actively stabilized on CO-associated bivalents in the same nucleus, which could result in redistribution of nucleoplasmic subunits and lowering of local protein concentrations below a threshold needed to reliably maintain SYP association on bivalents where stabilization has not occurred. Further, our analysis of SYP dynamics can likewise explain the even distribution of SYP proteins in nuclei where all chromosomes lack recombination sites. To reconcile this observation with a model wherein chromosomes lacking CO intermediates actively trigger desynapsis, Machinova et al. postulated that DSBs are required to activate the proposed surveillance mechanism. However, the observation is readily explained (without the need to invoke DSB-dependent activation of the surveillance system) by persistence of the more highly dynamic early pachytene state for all SCs, which would result in the SYP proteins remaining well distributed among them despite the fact that the SCs are not stabilized.

Second, we propose that CO-intermediate-triggered SC central region stabilization may be an integral feature of the CO regulation (aka “CO control”) system that operates to promote and ensure the formation of COs while at the same time limiting their numbers. Prior work showed that the *C. elegans* SYP proteins function both in promoting CO formation and inhibiting excess COs (22, 29, 30), leading Libuda *et al.* to propose that meiotic CO regulation may function as a self limiting system in which the SC or its components initially create an environment that promotes the formation of CO-eligible recombination intermediates, which in turn triggers a change in the state of the SC that antagonizes the formation of additional COs. We have now demonstrated that formation of CO intermediates is indeed coupled to a stabilization of the SC central region. We speculate that the stabilized SC central region may help both to a) facilitate correct maturation of CO intermediates at the sites that had been selected to become COs, and b) create a barrier to additional CO, *e.g.* by excluding DNA ends from engaging the homolog as a repair template and/or by promoting eviction of pro-CO proteins from recombination sites not selected to become COs.

Finally, we suggest that CO-intermediate-triggered SC stabilization likely plays a role in solidifying pairwise partnerships between homologous chromosomes. While the SC is required to stabilize associations between homologs during *C. elegans* meiosis (22), it is well established that formation of SCs between homologs in *C. elegans* does not depend on recombination (31). Indeed, the Caenorhabditis genus has lost an entire functional module of genes encoding conserved proteins needed to promote inter-homolog DNA strand invasion (DMC1/MND1/HOP2) in organisms where SC assembly is recombination-dependent (48). However, our recent work using polyploidy as a means to perturb the meiotic program revealed that recombination can nevertheless contribute to formation of stable pairwise interactions between homologs under conditions where three partners compete for synapsis (44). Our findings led us to propose a two-phase model for *C. elegans* synapsis involving a) an early phase in which synapsis interactions are driven by a recombination-independent homology assessment mechanism, and b) a late phase in which recombination promotes mature synapsis. The current work strongly reinforces this view. Although the SCs formed in the absence of recombination may appear “normal” (31), the assays used in the current work reveal that they do in fact differ from normal SCs. We speculate that the early, recombination-independent form of the *C. elegans* SC may function to mediate/maintain close homolog juxtaposition to facilitate productive engagement of the homologous chromosome by processed DSBs, whereas the late CO-intermediate-stabilized form may be analogous to the recombination-dependent SCs of budding yeast, Arabidopsis, and mammals.

## Materials and Methods

### Genetics

Unless otherwise specified, worms were cultured at 20°C under standard conditions (49). The following *C. elegans* strains were used:

- N2
- AV568: *meIs9[unc-119(+)Ppie-1::gfp::syp-3]*III; *unc-119(ed3)*III
- AV749: *syp-3(ok758)* I; *meIs9[unc-119(+)Ppie-1::gfp::syp-3]*
- AV779: *meIs9[unc-119(+)Ppie-1::gfp::syp-3]*III; *spo-11(me44) / nT1[qIs51]* IV,V
- AV794: *cosa-1(tm3298) / qC1[qIs26]* III; *meIs9[unc-119(+) Ppie-1::gfp::syp-3]*III
- AV899: *ieSi11 [cbunc-119+, Psyp-3::EmeraldGFP::syp-3]* II; *unc-119(ed3)* III; *ieSi21[sun-1::mRuby]* IV
- AV898: *syp-3(ok758)* I; *ieSi11 [cbunc-119+, Psyp-3::EmeraldGFP::syp-3]* II; *ieSi21[sun-1::mRuby]* IV
- AV900: *ieSi11 [cbunc-119+, Psyp-3::EmeraldGFP::syp-3]* II; *unc-119(ed3)* III *cosa-1(tm3298) / qC1[qIs26]* III; *ieSi21[sun-1::mRuby]* IV
- AV906: *ieSi11 [cbunc-119+, Psyp-3::EmeraldGFP::syp-3]* II; *unc-119(ed3)* III *spo-11(me44) ieSi21[sun-1::mRuby] / nT1[qIs51]* IV;V
- AV920: *plk-2(ok1936)* I*; ieSi11 [cbunc-119+, Psyp-3::EmeraldGFP::syp-3]* II; *ieSi21[sun-1::mRuby]* IV
- AV776: *spo-11(me44)* IV */ nT1[qIs51 let-?]* (IV;V)
- AV602: *spo-11(ok79)* IV / *nT1[qIs51 let-?]* (IV;V)
- CA1105: *dsb-1(tm5034)* IV / *nT1[unc-?(n754) let-?]* (IV;V)
- AV115: *msh-5(me23)* IV / *nT1[unc-?(n754) let-?]* (IV;V)
- AV596: *cosa-1(tm3298)* / *qC1[qIs26]* III
- AV872: *cosa-1(tm3298) / qC1[qIs26]* III; *spo-11(me44) IV / nT1[qIs51 let-?]* (IV;V)
- AV873: *+/szT1 I; spo-11(me44)/nT1[qIs51 let-?]* (IV;V); *dpy-8(e1321) unc-3(e151)* / *szT1[lon-2(e678)]* X
- AV831: *meIs8[unc-119(+); Ppie-1::gfp::cosa-1]* II; *cosa-1(tm3298)* III; *spo-11(me44)/nT1[qIs51 let-?]* IV;V
- AV883: *plk-2(ok1336) I; dsb-2(me96) meIs8[unc-119(+); Ppie-1::gfp::cosa-1]* II; *cosa-1(tm3298)* III
- CA1253: *syp-3 (ok758)* I; *ieSi11 [cbunc-119+, Psyp-3::EmeraldGFP::syp-3]* II; *unc-119(ed3)* III
- AV157: *spo-11(me44)/nT1[unc-?(n754)let-?qIs50] IV; nT1/ + V*
- AV818: *meIs8[unc-119(+); Ppie-1::gfp::cosa-1]* II; *cosa-1(tm3298)* III
- AV883: *plk-2(ok1336)* I; *meIs8[unc-119(+); Ppie-1::GFP::COSA-1]* II; *cosa-1(tm3298)* III
- AV917: *plk-2(me107[plk-2::HA])* I
- AV918: *dsb-2(me96) meIs8[unc-119(+); Ppie-1::gfp::cosa-1]* II; *cosa-1(tm3298)* III
- AV919: *plk-2(me107[plk-2::HA])* I; *dsb-2(me96)* II

A strain expressing PLK-2 with a C-terminal HA tag was generated by CRISPR/CAS9 genome engineering of the endogenous *plk-2* locus using the protocol of (50). A BamHI site was introduced as a spacer/screening sequence between the HA-tag and the last amino acid of PLK-2, resulting in following modification at the *plk-2* locus:

(*plk-2* to codon 632) GGATCCTACCCATACGACGTCCCAGACTACGCCTAA (*plk-2* 3’UTR).

### Cytological analysis of whole-mount gonads

All analyses were performed on 20-24h post-L4 adults. For immunofluorescence experiments on whole-mount gonads, dissection of gonads, fixation, immunostaining and DAPI counterstaining were performed as in (51). The following primary antibodies were used: chicken anti-HTP-3 (1:250 (MacQueen et al. 2005)), rabbit anti-GFP 12)), guinea-pig anti-SYP-1 (1:200 (22), rabbit anti-MSH-5 12)). Secondary antibodies were Alexa Fluor 488, 555 and 647-conjugated goat antibodies directed against the appropriate species (1:400, Life Technologies). Images in Figs 1, 4, 5 and 7C were collected as Z-stacks (at 0.2μm intervals) using a 100x NA 1.40 objective on a DeltaVison OMX Blaze microscopy system (except for the 7C *plk-2* single mutant, which was acquired using a 63X NA 1.40 objective), deconvolved and corrected for registration using SoftWoRx. Multiple overlapping fields covering the whole length of the gonad were acquired for each specimen, and gonads were assembled using the “Grid/Collection” plugin (32) on Fiji. Final assembly of 2D maximum or sum intensity projections was performed using Fiji (52), with adjustments of brightness and/or contrast made in Fiji or Adobe Photoshop.

### Quantitation of EmGFP::SYP-3 fluorescence in gonads of live worms

Worms were prepared and imaged using the OMX microscope as described below for FRAP analysis (except that serotonin was not included); slides were imaged within 15 minutes after mounting. For Figs 1B and S1 (increase of SYP-3 during pachytene progression), we generated a sum intensity projection of all the stacks containing the nuclei of the top layer of the gonad; for each nucleus fully contained within this projection, we calculated the average fluorescence intensity of that nucleus as the total fluorescence in the smallest circle containing the nucleus, divided by the area of the circle.

### Quantitation of nuclei exhibiting uneven SYP-1 distribution

For Fig 4, pachytene nuclei were manually segmented on maximum intensity projections and scored for the presence of an uneven distribution of SYP-1 among their SCs. Nuclei in the second half of the pachytene region were assessed for presence or absence of this feature; the pachytene region was defined as spanning the distance between the most distal nucleus with six SC stretches and unclustered chromosomes to the last row containing multiple nuclei.

For Fig S6, we quantified the number of chromosome pairs that exhibited SYP-1 enrichment (within each scored nucleus); for this feature, we scored nuclei in the last quarter of the pachytene region, where the distinction between SYP-1-enriched chromosomes and SYP-1-depeleted chromosomes was most pronounced.

### Detergent lysis, nuclear spreading and immunofluorescence of *C. elegans* meiocytes

The protocol used for detergent lysis and spreading of *C. elegans* germ cell nuclei (Figs S2-S4, 5 and 6) is an adapted version of the protocol used in (53) for *S. cerevisiae*. The gonads of 20-100 adult worms were dissected in 5μl dissection solution (see note A) on a 22×40mm coverslip (thickness #1.5, as required for the OMX microscope). 50μl of spreading solution (see note B) was added and gonads were immediately distributed over the whole coverslip using a glass rod or a pipette tip. Coverslips were left to dry overnight at room temperature or for two hours at 37°C, washed for 20 minutes in methanol at -20°C and rehydrated by washing 3 times for 5 minutes in PBS-T. A 20 minute blocking in 1% w/v BSA in PBS-T at room temperature was followed by overnight incubation with primary antibodies at 4°C (antibodies diluted in: 1% w/v BSA in PBS-T supplied with 0.05% w/v Sodium azide). Coverslips were washed 3 times for 5 minutes in PBS-T before secondary antibody incubation for 2 hours at room temperature. After PBS-T washes, the nuclei were immersed in Vectashield and the coverslip was mounted on a slide and sealed with nail polish. The following primary antibodies were used: chicken anti-HTP-3 (1:500 (42)), rabbit anti-GFP (1:750 (12)), guinea-pig anti-SYP-1 (1:200 (22)), rabbit anti-MSH-5 (1:10000 (SDIX)) and mouse anti-HA (Covance)

### Note A

Dissection solution: The baseline dissection solution is: 0.1% v/v Tween-20 in water. This leads to the strongest wash-out of non-chromatin bound factors, but also leaves only a few gonads on the slide in which the temporal/spatial organization of the germline nuclei remains unperturbed. Increasing salt concentration by adding PBS, Dulbecco’s Modified Eagle’s Medium (DMEM, Sigma-Aldrich, D6546) or similar, results in more intact germline organization, but less complete washout of non-chromatin bound factors.

### Note B

Spreading solution (for one coverslip, 50μl): 32μl of Fixative (4% w/v Paraformaldehyde and 3.2-3.6% w/v Sucrose in water), 16μl of Lipsol solution (1% v/v/ Lipsol (obtained from Josef Lodl) in water), 2μl of Sarcosyl solution (1% w/v of Sarcosyl in water). Increasing Sarcosyl concentration increases wash out of non-chromatin bound factors.

### FRAP analyses

### Strains used

FRAP experiments using the Leica SP2 confocal microscope were conducted using strains derived from AV568, which contains *meIs9*, a transgene insertion generated by microparticle bombardment (54). GFP::SYP-3 expressed from this transgene in the presence of a wild-type *syp-3* allele exhibits robust localization identical to endogenous SYP proteins, and AV568 germ lines exhibit normal meiosis. However, *meIs9* only partially rescues the phenotypes of a *syp-3* null mutant, apparently enabling substantial chiasma formation without restoring normal chromosome segregation (Fig S7), and we subsequently discovered that a missense mutation causing a K159R amino acid substitution had been introduced during generation of the transgenic line. Thus, we also conducted a full set of FRAP analyses using strains containing the *ieSi11* transgene (55), which was generated by the MosSCI method; *ieSi11* provides much better, albeit still incomplete, rescue of the *syp-3* null phenotype (55)( Fig S7), so experiments using *ieSi11* were also conducted in strains carrying the wild-type *syp-3* allele.

### Sample preparation and recovery

L4 hermaphrodites were picked to a fresh plate and upshifted to 25°C for approximately 24 hr prior to each FRAP experiment. Worms were placed on slides with 7% agarose pads, and a mixture of 1 μl of silica microspheres (Bangs Laboratories, Cat# SS02N/11167) and 0.4-0.6 μl of anesthetic solution (0.2% tricaine + 0.02% tetramisole) was added to the pad. For experiments done on the Deltavision OMX microscope (see below), 0.5 μl of serotonin solution (55) (final concentration of 25mM prepared from Serotonin Hydrochloride salt, Sigma) was added together with the silica beads and anesthetic solution. 5-7 worms were then mounted with a small amount of bacteria (in order to maintain healthy conditions during imaging). The coverslip (22X40mm-1.5) was sealed to the slide using a thin line of petroleum jelly applied by syringe around the agarose pad; mild pressure was applied to achieve an even spread of liquid on the pad and to ensure efficient immobilization of the worms. After imaging, the cover slip was removed and worms were transferred to a seeded NGM plate and allowed to recover for 1-2 days; worms were monitored for movement and egg laying to verify their recovery.

### FRAP experiments on Leica SP2 Confocal

A Leica SP2 laser scanning confocal microscope was used to perform FRAP experiments. Images were acquired using a 63X oil HCX PL APO objective (N. A 1.32) (Leica Microsystems, Heidelberg, Germany). Worms were located using wide field illumination, and confocal imaging was carried out using a 20mW Argon laser for excitation of GFP at 488nm. The Leica confocal software (v. 2.5 build 1347) was used to control all scanning and acquisition parameters. Images were acquired using an 8X zoom at a resolution of 512×512 pixel dimensions as z stacks at 0.24 μm intervals. The laser power used for the bleach step was approximately 7x higher than the power used for prebleach and postbleach imaging. The automated FRAP module built into the Leica confocal software was used to perform the initial steps of the FRAP experiment, but the automated module could not be used for the entire experiment because nuclei continued to progress proximally through the gonad during the course of the experiment, resulting in substantial shifts in position along the x and y dimensions (lateral displacement on the order of 5-10 μm over the course of an experiment), as well as some shift in z, thus requiring manual readjustment of the imaging field.

For each experiment, a field of pachytene nuclei was selected and a range of z stacks encompassing the nuclei was set. In the FRAP module, a snapshot of this field was acquired. Regions of interest (ROIs) were drawn manually in the field, selecting approximately 30-50% of the area of three nuclei (rectangular ROIs) to partially bleach, and the entire area of one nucleus (elliptical ROI) to fully bleach. The FRAP module was programmed to acquire one prebleach Z stack, perform the bleach step through every step of the Z series (spending ~0.8 s per z slice), and then immediately acquire one postbleach Z stack. Imaging was continued for a period of 1 hour, at time points as depicted in Fig 3A-B. At the beginning and end of each experiment, an image of the whole gonad was acquired (at 2x zoom) in order to determine the position of the bleached nuclei within the pachytene zone of the gonad. Gut autofluorescence was also checked at the end of the imaging period to verify the health and viability of the worm, and the worm was recovered as described above.

All images analyzed were sum intensity projections of Z stacks generated using ImageJ. This approach was taken as it is robust in the face of the substantial movement of nuclei in the x, y and z dimensions (discussed above), which would introduce considerable noise into measurements taken for individual z-sections. Excess z-sections above and below the nuclei were excluded and the number of z-sections per projection was kept constant within an experiment. Circular segment ROIs were drawn in ImageJ using the ‘oval’ and ‘rectangular’ tools to measure fluorescence intensity within the bleached and unbleached portions of 3 partially bleached nuclei; the chord separating the two circular segments was deliberately drawn well within the portion of the nucleus that had been bleached, avoiding the boundary between the bleached and unbleached portions, to ensure that the ROIs used to assess recovery of bleached SCs only contained chromosome segments that had been bleached. Fluorescence intensities were also measured for 1 fully bleached nucleus, and 1 unbleached reference nucleus; 3 smaller ROIs were drawn in portions of the image outside the nuclei as a reference for subtracting background fluorescence. All ROIs drawn were added to the ROI manager tool. The area and the total fluorescence intensity within each ROI was then determined for each time point using the ‘measure’ function in ROI manager. Positions of ROIs were adjusted to correct for the forward progression of the nuclei within the gonad. Patterns of GFP fluorescence within nuclei were also examined visually, both in projections and in image stacks. In cases where substantial movement of chromosomes or rotation of a nucleus altered the pattern of the GFP in the nucleus such that fluorescent SC segments that were originally within an “unbleached” ROI had moved into the “bleached” ROI, such nuclei were excluded from quantitation.

For each nucleus analyzed, the extent of recovery at post-bleach time points was quantified by calculating the ratio of fluorescence signal within an ROI containing only the bleached portion of the nucleus to the fluorescence signal for that whole nucleus, normalized to the same ratio observed prior to the bleach; prior to calculating these ratios, fluorescence values for each ROI were corrected for background fluorescence. Statistical analyses were conducted using GraphPad Prism software.

### FRAP experiments on DeltaVision OMX

A DeltaVision OMX Blaze microscopy system (Applied Precision/ GE healthcare) was used to perform FRAP experiments in conventional widefield mode. Images were acquired using a 100X oil objective (U-PLANSApo, N.A 1.42). At the beginning and end of each experiment, the spiral mosaic function was used to generate a composite image of the gonad in order to determine the position of the bleached nuclei within the pachytene zone of the gonad. Imaging was carried out by means of EMCCD cameras (Photometrics Evolve 512) and bleaching was performed using a 100mW 488nm laser for excitation of GFP. The DeltaVision OMX software (version 3.60.7848.0) was used to control all scanning and acquisition parameters. Once the z range was set, images were acquired at a resolution of 256X×256 pixel dimensions as z stacks at 0.24 μm intervals. The EMCCD camera was set to image at 10% transmission and 100ms exposure (adjusted depending on fluorescence intensity) in ‘conventional’ mode. The bleach step was carried out using ‘photo kinetic’ mode, at 10% laser power, for a duration of 0.2s on a chosen slice. In contrast to the Leica SP2 confocal system where the bleach could be performed through every step in a z-series, the FRAP module in the OMX widefield system uses a photobleaching beam that is directed to a single selected focal plane. For these experiments, bleaching was focused on a selected plane in the top half of the nucleus in which multiple stretches of SC were visible.

For the experiments presented in Figs 2D-F, 3A and 3C, a prebleach z-series was first acquired in both green (EmGFP::SYP-3) and red (SUN-1::mRuby, (55)) channels spanning full nuclei in the chosen field. A prebleach image of the SUN-1::mRuby was taken to record the meiotic state of each nucleus in the field. A focal plane with a large number of SYP stretches was then located in the green channel, and an image of the nuclei was acquired. Rectangular ROIs between 1 − 3 μm^2^ were drawn on 4−5 nuclei in order to bleach limited stretches of fluorescence. The bleach step was performed on this plane. A post-bleach z-series image was then quickly acquired with the original range used in the prebleach z stack. Images were acquired for a period of 30 minutes post bleach, with manual adjustments of the position of the field whenever required. At the end of the experiment, a second spiral mosaic image of the whole gonad was acquired to confirm the position of bleached nuclei. Images were deconvolved using DeltaVision SoftWorx software (version 6.1.1).

All images analyzed were sum intensity projections of deconvolved z stacks containing the top half of each nucleus analyzed, generated using Fiji ImageJ. Partial projections were used to avoid overlaying SYP stretches from the top and bottom halves of the nucleus, thereby enabling unambiguous visualization of the bleached SYP stretches. (Full image stacks were also examined to verify that observed loss and restoration of pattern elements reflected bleaching and subsequent recovery rather than nuclear or chromosomal movement.) Excess z-sections above the nuclei were excluded and the number of z-sections per projection was kept constant within an experiment. To enable unbiased scoring of recovery in individual nuclei, each bleached nucleus scored was cropped from the corresponding prebleach, post-bleach, 5min, 20min and 30min sum intensity projections and pasted in a panel as shown in Figs 2D and 3A. Such panels were prepared for 3 stages of pachytene nuclei (early, mid, late) from 3 genotypes (WT, *spo-11, cosa-1*). The same approach was used for comparing late pachytene nuclei from WT and the *plk-2* mutant in Fig 7.

The combined use of small bleach ROIs and serotonin in the mounting preparation helped to preserve normal physiological movement of chromosomes within pachytene nuclei, resulting in modest changes in the relative shape or position of SYP stretches within scored nuclei. Thus, we devised a double-blind scoring system in which 1) stage and genotype identifiers of panels were removed and 2) two independent investigators not involved in the acquisition of the images or the preparation of panels scored each of the panels for the occurrence of strong fluorescence redistribution by the 30min time point. The graphs presented in Fig 2E, 3C and 7B plot the percentage of nuclei that were assessed concordantly as exhibiting strong recovery (defined as apparent evening-out of fluorescence between bleached and unbleached SC segments) by the two independent scorings in this double-blind scheme. Numbers of nuclei scored for Figs 2 and 3 were: (WT early, n = 28; WT mid, n = 14; WT late, n = 27; *spo-11* early, n = 19; *spo-11* mid, n = 30; *spo-11* late, n = 28; *cosa-1* early, n = 22; *cosa-1* mid, n = 22; *cosa-1* late, n = 21). Numbers of nuclei scored for Fig 7 were: (WT late, n = 24; *plk-2* late, n = 23). Statistical analyses were conducted using GraphPad Prism or GraphPad QuickCalcs software.

## Acknowledgements

We thank the *Caenorhabditis* Genetics Center (funded by NIH Office of Research Infrastructure Programs P40 OD010440) for strains and A. Dernburg for strains and antibodies. We thank Grace Chen and Marc Presler for early work that helped to establish experimental conditions for the FRAP analyses. This work was supported by an FWF Erwin Schrödinger Fellowship (J-3676) to A. W. and by NIH grant R01GM53804 and American Cancer Society Research Professor Award RP-15-209-01-DDC to A.M.V. The project was also supported, in part, by Award Number 1S10OD01227601 from the National Center for Research Resources (NCRR). Its contents are solely the responsibility of the authors and do not necessarily represent the official views of the NCRR or the National Institutes of Health.

## Supporting information captions

**Figure S1:**
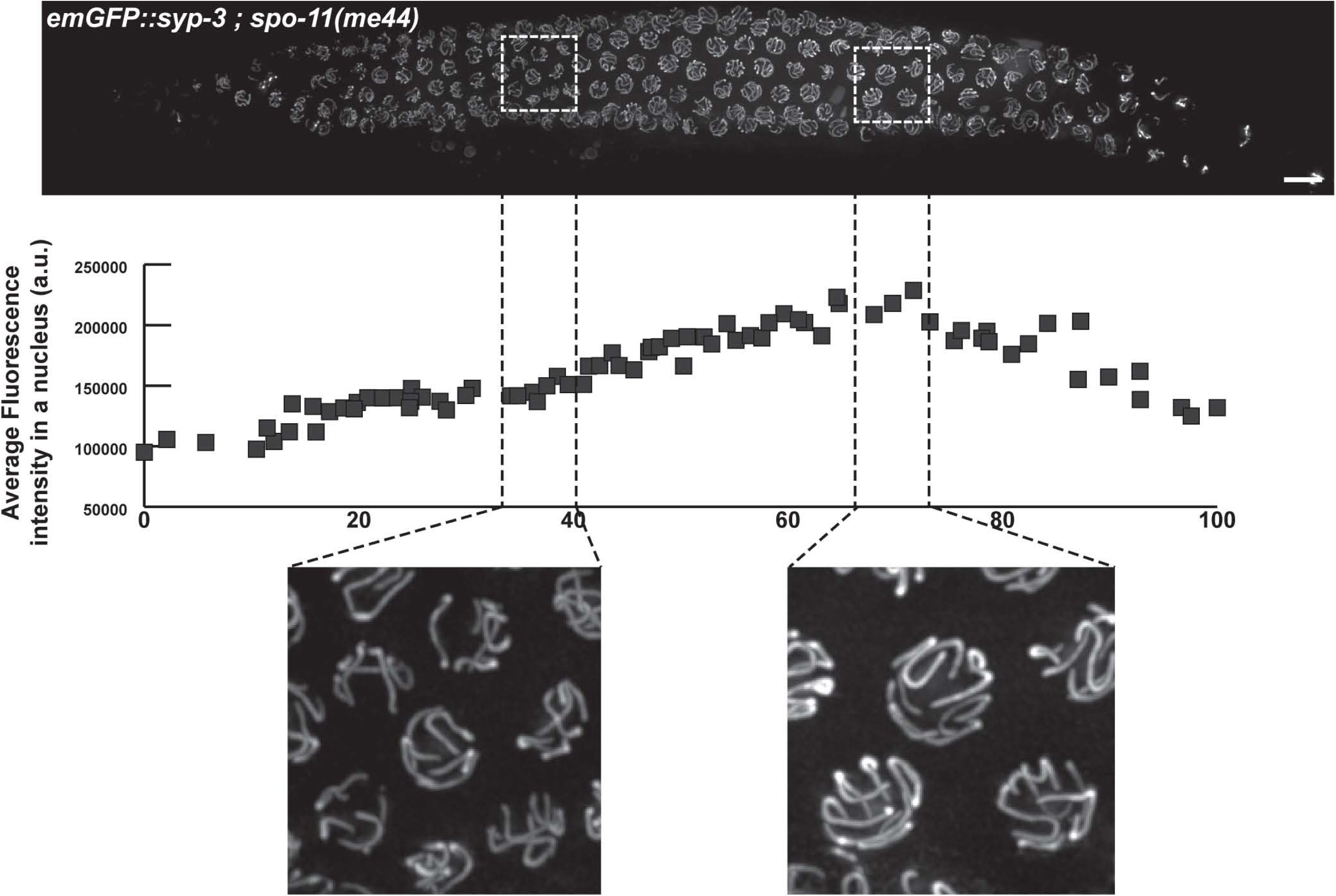
Levels of SC central region component SYP-3 increase during pachytene progression in *spo-11* mutant meiocytes. Detection of EmGFP::SYP-3 in the gonad of a live *spo-11* mutant worm (top). The graph (middle) represents the evolution of the average fluorescence intensity of a nucleus as a function of its position in the gonad (see Materials and Methods). The increase in GFP::SYP-3 signal upon pachytene progression is illustrated in higher magnification fields of early and late pachytene nuclei (bottom). Images are maximum intensity projections of 3D data stacks encompassing whole nuclei. Scale bar represents 10μm.

**Figure S2:**
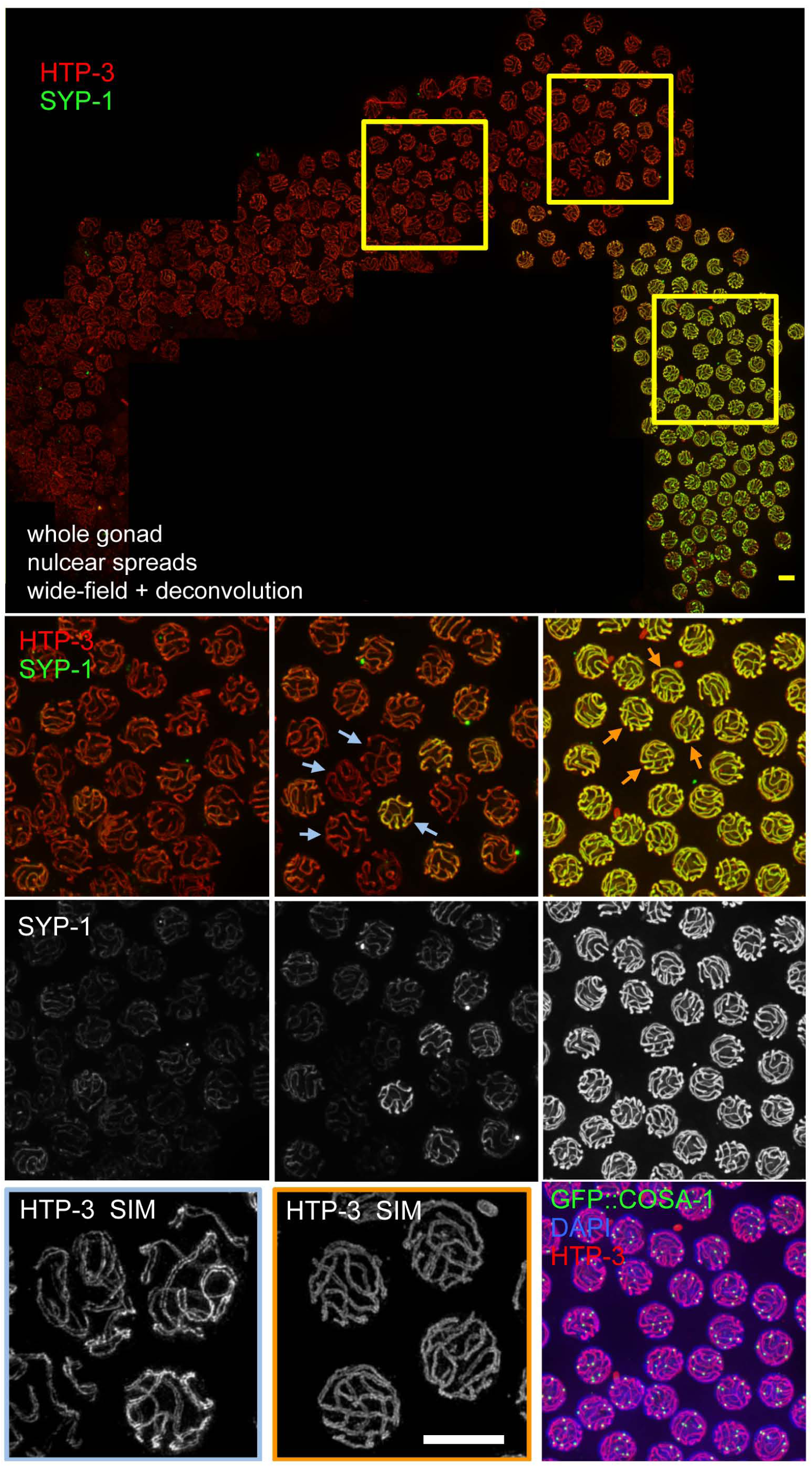
Differential sensitivity of early and late pachytene nuclei to removal of SYP-1 by detergent-based lysis. Immunofluorescence images of germ cell nuclei from a *C. elegans* gonad subjected to detergent-based lysis and partial nuclear spreading using a procedure where the temporal-spatial organization of the gonad can be substantially preserved. Fields of nuclei from early, middle (early-to-late pachytene transition), and late pachytene regions of the gonad, showing that HTP-3 (top row of insets) is retained on chromosome axes following this procedure, whereas SC central region protein SYP-1 (middle row) is largely removed from early pachytene nuclei, yet strongly retained in late pachytene nuclei. The center SYP-1 panels show that adjacent nuclei in the early-to-late pachytene transition region can exhibit different behavior regarding retention/removal of SYP-1. Bottom right inset: late pachytene nuclei with strong SYP-1 retention also exhibit another late pachytene marker, namely the presence of 6 bright GFP::COSA-1 foci marking the sites of CO-designated recombination events. As depicted in the accompanying Fig S3, the transition to the state where SYP-1 is strongly retained coincides with acquisition of this late pachytene stage marker. Bottom left and center: 3D-SIM images of HTP-3 localization in nuclei from the early-to-late pachytene transition (left, blue outline) and late pachytene (center, orange outline) regions of the gonad; the positions of the depicted nuclei are indicated by blue or orange arrows in the top row of insets. The nuclei where SYP-1 was depleted exhibit wider separation between aligned HTP-3-marked axes than adjacent nuclei where SYP-1 was retained. Images are maximum intensity projections of 3D data stacks; scale bars represent 5 μm

**Figure S3:**
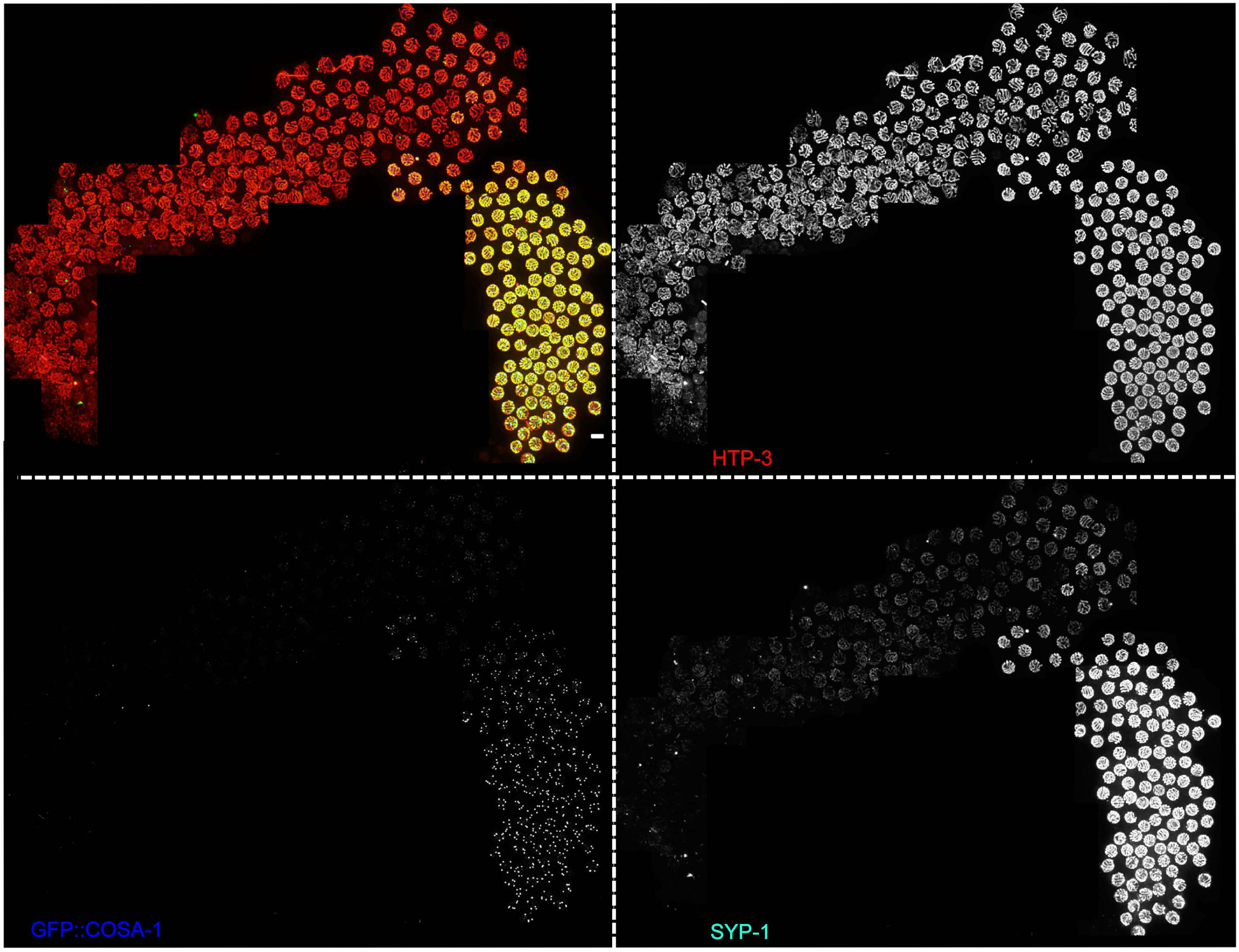
Differential sensitivity of early and late pachytene nuclei to removal of SYP-1 by detergent-based lysis. Images of the same spread gonad depicted in Fig S1, showing HTP-3, SYP-1 and GFP::COSA-1 channels separately to facilitate visualization of the transition from the early to late pachytene state, as indicated by the presence of 6 bright GFP::COSA-1 foci.

**Figure S4:**
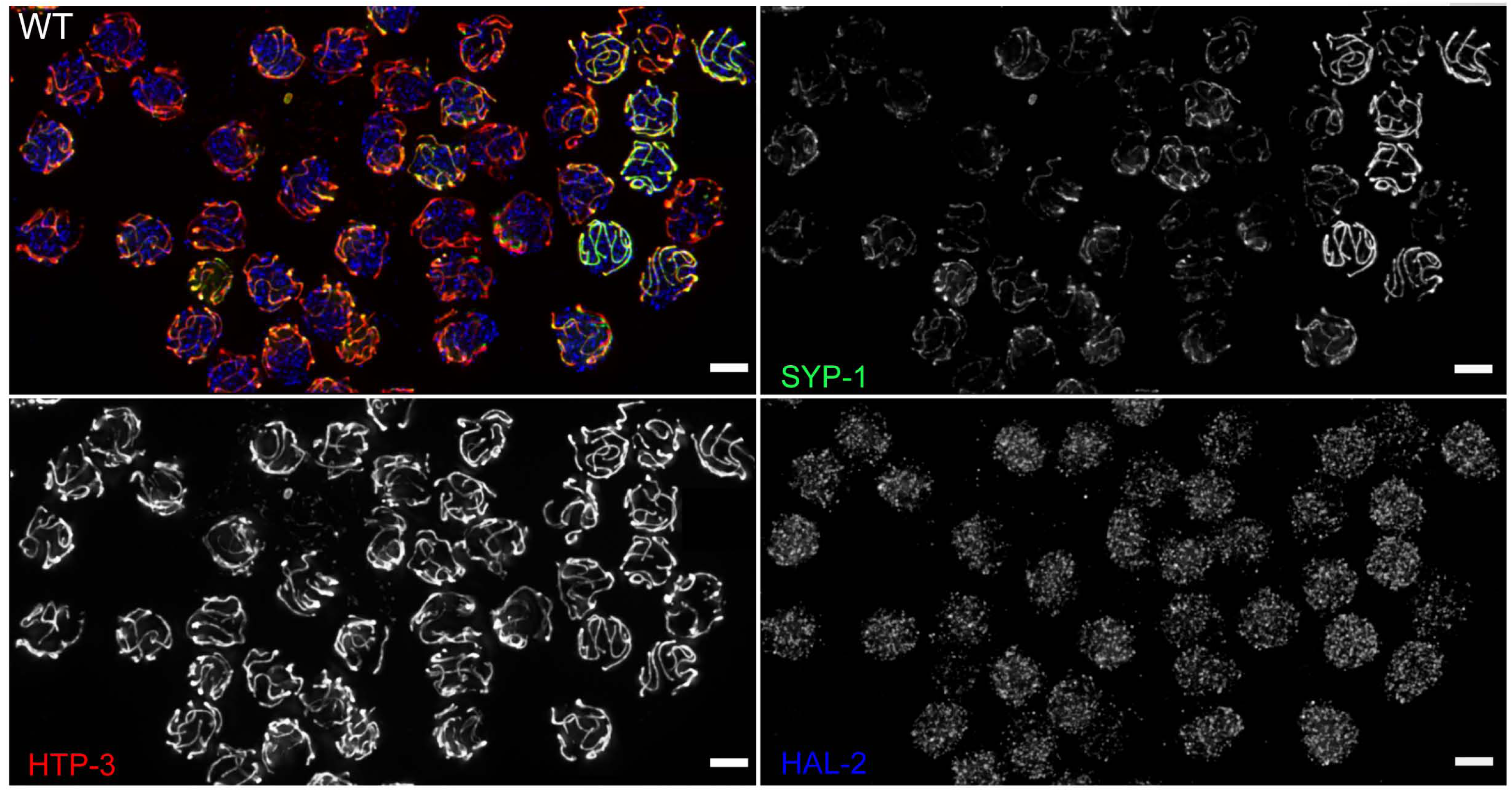
Differential sensitivity of early and late pachytene nuclei to removal of SYP-1 is not solely a consequence of differential susceptibility to nuclear disruption and spreading. Representative field of nuclei from the early-to-late pachytene transition region of a WT gonad subjected to detergent-based lysis and immunostained to detect localization of HTP-3 (axis), SYP-1 (SC central region) and HAL-2 (a nucleoplasmic protein). As we have observed that nuclei in earlier stages of prophase are in general more susceptible than late prophase nuclei to disruption and spreading by detergent-based lysis procedures, we wanted to rule out the possibility that the higher sensitivity of SYP-1 to removal at earlier stages might be explained solely as a secondary consequence of this higher sensitivity to nuclear disruption. To this end, we carried out simultaneous immunoloclaization of SYP-1 and HAL-2, a nuclear protein that is concentrated in the nucleoplasm but is not associated with chromatin. As HAL-2 is largely, but not completely, released from nuclei when they are partially disrupted by detergent lysis, the amount of residual HAL-2 detected in nuclei can serve as a proxy for the degree of nuclear disruption. In the nuclei depicted here, we did not observe a correlation between the amount of residual HAL-2 and SYP-1 after detergent lysis, indicating that the differential sensitivity of early and late pachytene nuclei to SYP-1 removal can be observed even when the overall degree of nuclear disruption is comparable. Thus, the higher sensitivity of SYP proteins to removal at earlier stages is not solely a secondary consequence of earlier nuclei being more susceptible to mechanical disruption, but rather implies a state change in the SCs themselves. Scale bar, 5μm.

**Figure S5:**
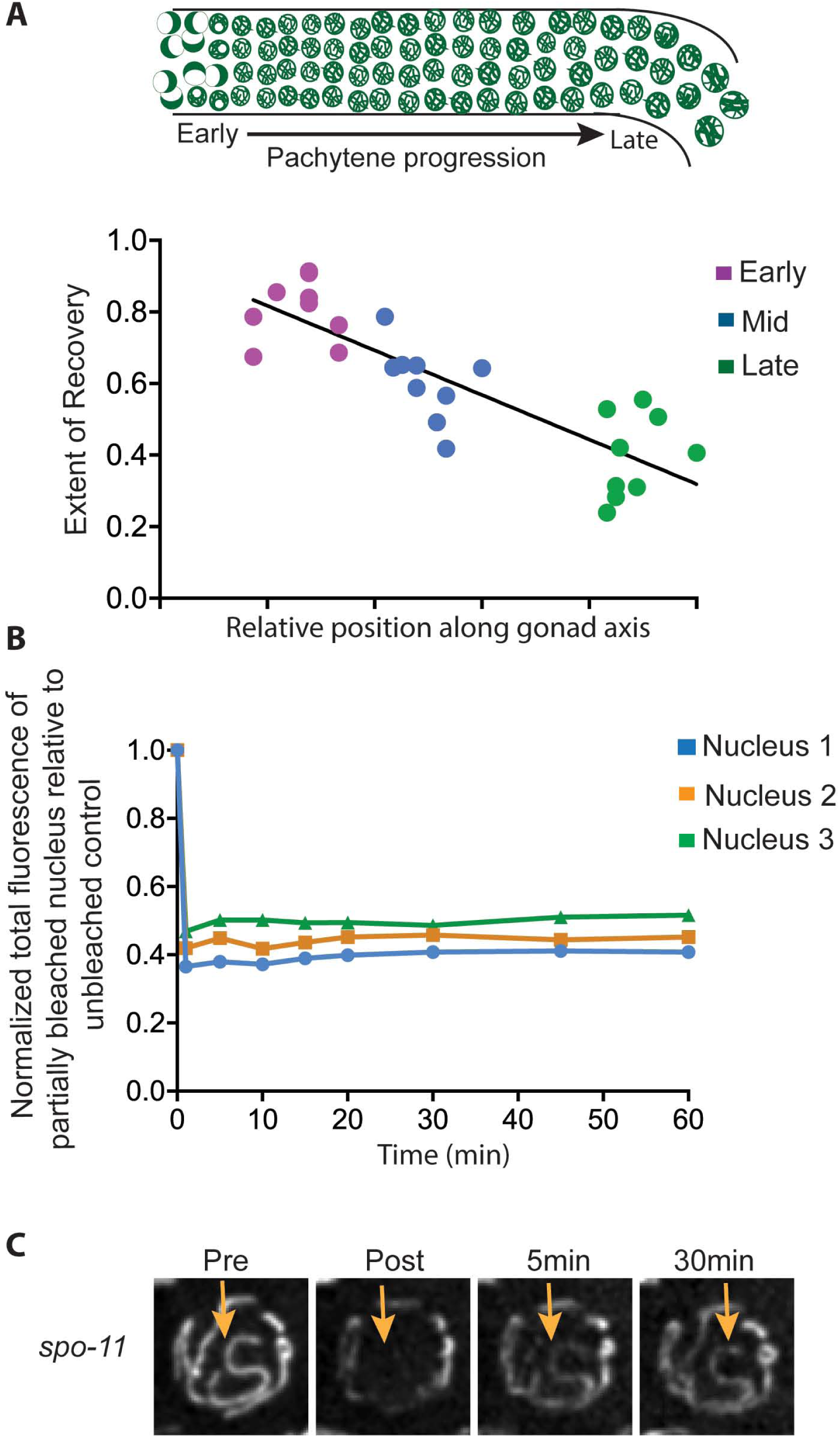
Features of SC dynamics revealed by FRAP analyses. **A.** Alternative representation of the quantitation of the confocal FRAP experiments presented in Fig 3B. The plateau values for the “extent of fluorescence recovery” curves for each nucleus were plotted against the position of that nucleus along the gonad axis. Relative position along the gonad axis was calculated by dividing the row number of the nucleus by the total number of pachytene rows in that gonad. Slope of best fit line = -0.62 (R^2^ = 0.71), reflecting a reduction in GFP::SYP-3 dynamics as nuclei progress through the pachytene stage. **B.** Graph plotting the total fluorescence in representative partially bleached nuclei (relative to an unbleached reference nucleus in the same field) from a FRAP experiment analyzing mid-pachytene nuclei in wild-type worms. Normalized total fluorescence in each partially bleached nucleus remains nearly constant during the recovery time course when compared to an unbleached reference nucleus. **C.** Similar to Fig 3E: panel of EmGFP::SYP-3 images of a *spo-11* mutant nucleus showing FRAP of an SC that had been bleached along its entire length (arrow).

**Figure S6:**
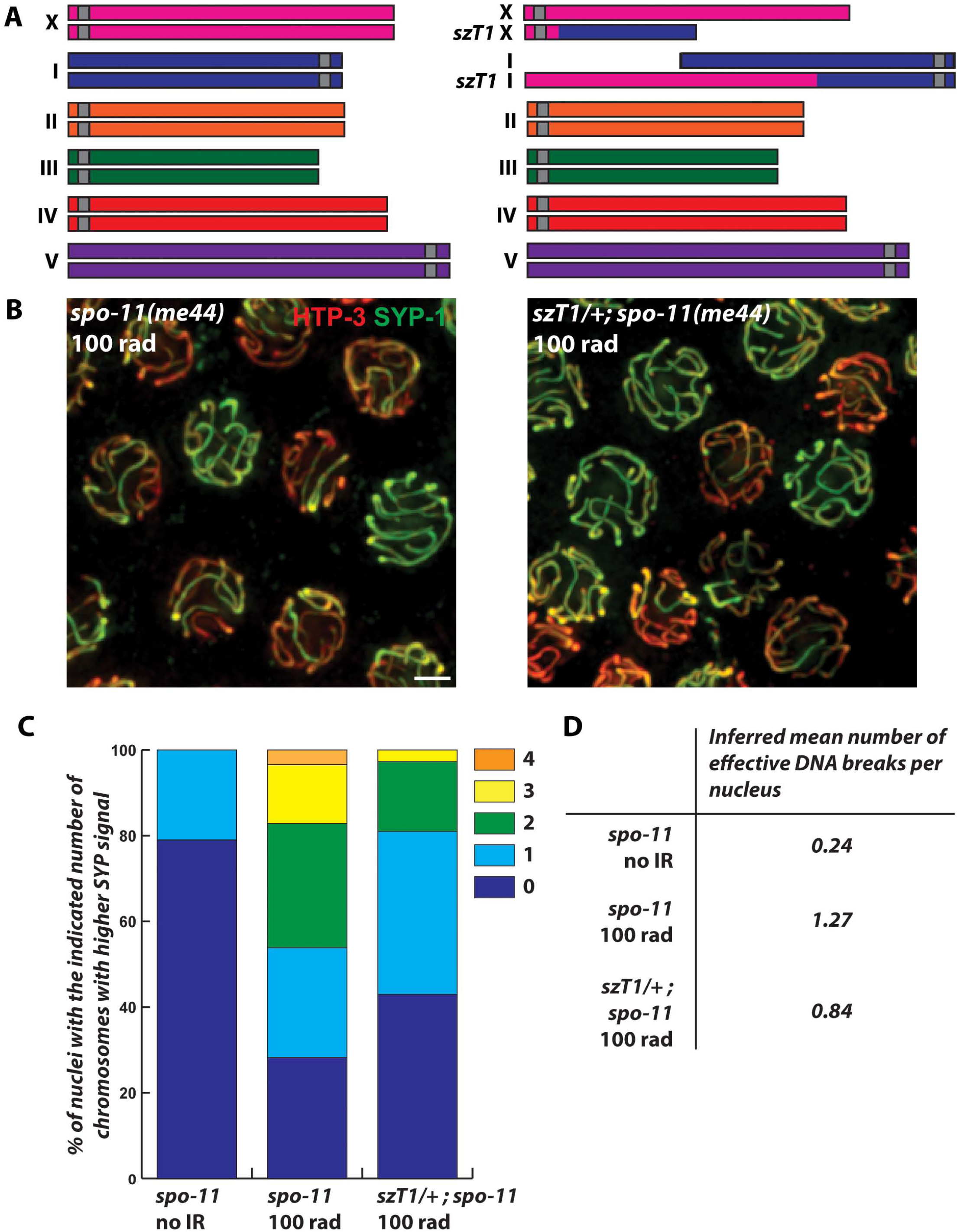
Irradiation-induced breaks can induce enrichment of SC central region components on some bivalents by a process dependent on homology between the synapsed partners. **A.** Schematic representation of synaptic configurations in *spo-11* mutant meiocytes (left) and *szT1* heterozygous *spo-11* mutant meiocytes (right). The *szT1* rearrangement is a reciprocal translocation between chromosomes *I* and *X,* and therefore in *szT1* heterozygous meiocytes, one third to one quarter of the genome is synapsed non-homologously. **B.** Representative fields of whole-mount late pachytene nuclei from *spo-11* (left) and *szT1* heterozygous *spo-11* (right), stained for HTP-3 and SYP-1 eight hours after exposure to 100 rad of γ-irradiation, showing an increased incidence of nuclei with uneven SYP-1 distribution. Scale bars represent 2μm. **C.**. Quantitation of the proportions of nuclei in the last quarter of the pachytene region with the indicated number of chromosome pairs exhibiting SYP-1 enrichment in gonads of untreated and irradiated *spo-11* worms and irradiated *spo-11* worms heterozygous for *szT1* (Of note, the samples analyzed for unirradiated *spo-11* are the same as in Fig 5, but were reanalyzed to focus exclusively on nuclei in the last quarter of pachytene, where SYP-1 enrichment is more pronounced, enabling quantitation of numbers of SYP-1-enriched chromosome pairs within the subset of nuclei exhibiting uneven SYP-1 distribution.) **D.** We estimated the mean number of DNA breaks per nucleus for each genotype and irradiation dose analyzed in panel C, based on a) previous work showing that in *C. elegans* meiocytes lacking SPO-11 induced DSBs, a single DSB can be efficiently converted into a CO recombination intermediate, and b) our hypothesis that enrichment of SYP-1 on a subset of chromosomes in late pachytene nuclei might be triggered by the formation of recombination intermediates. Thus, the mean number of DNA breaks that were effective in eliciting SYP-1 enrichment was inferred from the proportion of nuclei with even SYP-1 distribution (dark blue in panel C, = zero effective breaks), assuming a Poisson distribution. The magnitude of the reduction in inferred mean number of effective DNA breaks induced by IR in *szT1/+; spo-11* vs. *spo-11* is comparable to the fraction of the genome that is engaged in heterosynapsis in *szT1* heterozygotes.

**Figure S7:**
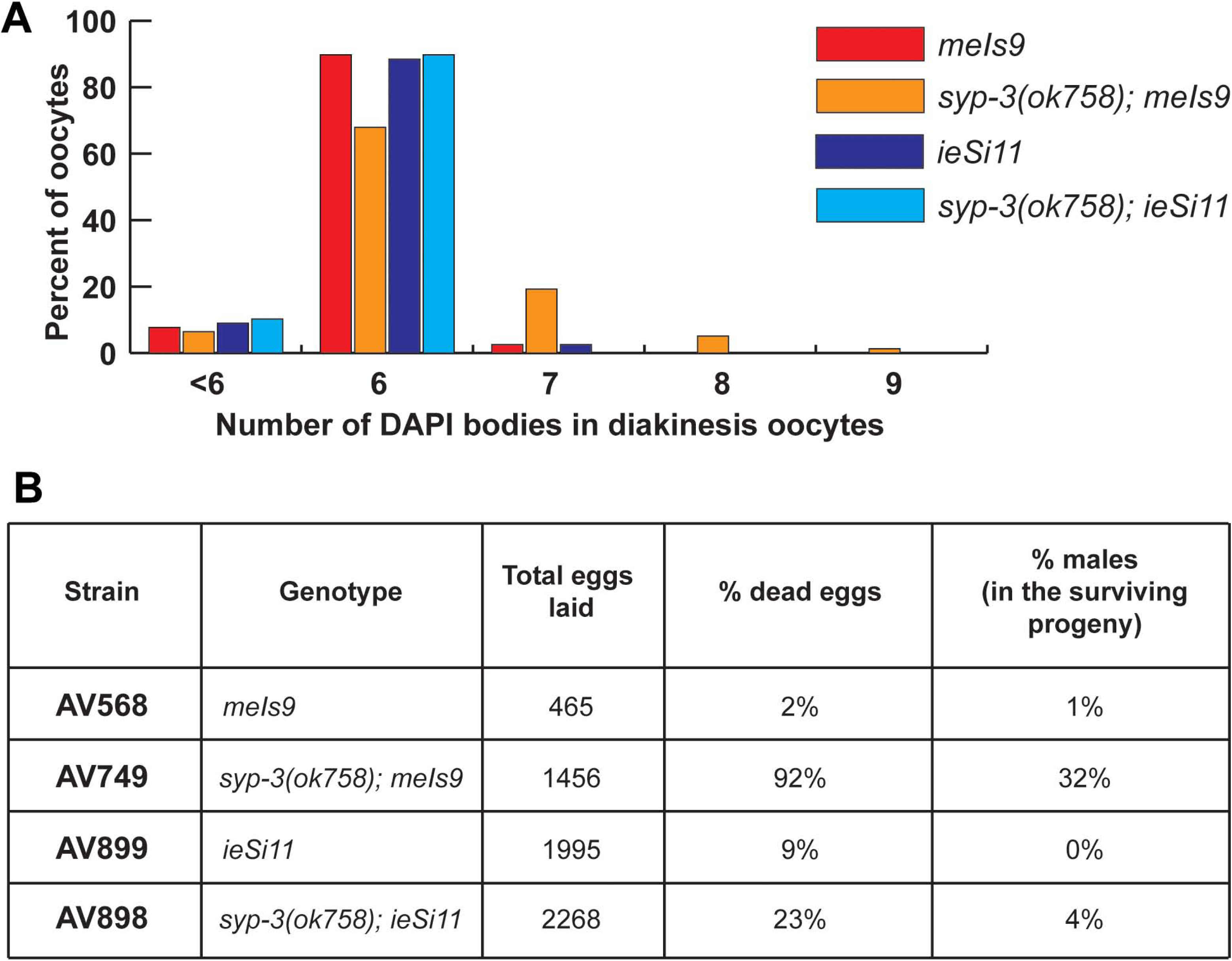
Characterization of strains carrying the *gfp::syp-3* transgenes used for FRAP analyses. **A.** Quantitation of DAPI bodies in oocytes at diakinesis, the last stage of meioticprophase. Wild type oocytes have 6 DAPI bodies, corresponding to six pairs of homologous chromosomes connected by chiasmata; observation of 7-9 DAPI bodies reflects a failure to form chiasmata between a subset of chromosome pairs. 78 nuclei diakinesis nuclei were scored for each genotype. **B.** Quantitation of percent males among surviving progeny, which reflects mis-segregation of X chromosomes, and inviability of embryos, which in the context of meiotic mutants can reflect mis-segregation of autosomes and/or defective repair of meiotic DSBs.

## References

1. Loidl J. Conservation and Variability of Meiosis Across the Eukaryotes. Annual review of genetics. 2016.

2. Muller HJ. The Mechanism of Crossing-Over. II. IV. The manner of occurrence of crossing-over. The American Naturalist. 1916;50(593):284–305.

3. Bergerat A, de Massy B, Gadelle D, Varoutas PC, Nicolas A, Forterre P. An atypical topoisomerase II from Archaea with implications for meiotic recombination. Nature. 1997;386(6623):414–7.

4. Keeney S, Giroux CN, Kleckner N. Meiosis-specific DNA double-strand breaks are catalyzed by Spo11, a member of a widely conserved protein family. Cell. 1997; 88(3):375–84.

5. Sun H, Treco D, Schultes NP, Szostak JW. Double-strand breaks at an initiation site for meiotic gene conversion. Nature. 1989;338 (6210): 87–90.

6. Szostak JW, Orr-Weaver TL, Rothstein RJ, Stahl FW. The double-strand-break repair model for recombination. Cell. 1983;33 (1): 25–35.

7. Hollingsworth NM, Ponte L, Halsey C. MSH5, a novel MutS homolog, facilitates meiotic reciprocal recombination between homologs in Saccharomyces cerevisiae but not mismatch repair. Genes & development. 1995;9 (14): 1728–39.

8. Kelly KO, Dernburg AF, Stanfield GM, Villeneuve AM. Caenorhabditis elegans msh-5 is required for both normal and radiation-induced meiotic crossing over but not for completion of meiosis. Genetics. 2000; 156 (2): 617–30.

9. Ross-Macdonald P, Roeder GS. Mutation of a meiosis-specific MutS homolog decreases crossing over but not mismatch correction. Cell. 1994;79 (6): 1069–80.

10. Zalevsky J, MacQueen AJ, Duffy JB, Kemphues KJ, Villeneuve AM. Crossing over during Caenorhabditis elegans meiosis requires a conserved MutS-based pathway that is partially dispensable in budding yeast. Genetics. 1999;153 (3): 1271–83.

11. Holloway JK, Sun X, Yokoo R, Villeneuve AM, Cohen PE. Mammalian CNTD1 is critical for meiotic crossover maturation and deselection of excess precrossover sites. The Journal of cell biology. 2014;205(5): 633–41.

12. Yokoo R, Zawadzki KA, Nabeshima K, Drake M, Arur S, Villeneuve AM. COSA-1 reveals robust homeostasis and separable licensing and reinforcement steps governing meiotic crossovers. Cell. 2012; 149 (1): 75–87.

13. Fawcett DW. The fine structure of chromosomes in the meiotic prophase of vertebrate spermatocytes. The Journal of biophysical and biochemical cytology. 1956;2(4): 403–6.

14. Moses MJ. Chromosomal structures in crayfish spermatocytes. The Journal of biophysical and biochemical cytology. 1956; 2 (2): 215–8.

15. Schild-Prufert K, Saito TT, Smolikov S, Gu Y, Hincapie M, Hill DE, et al. Organization of the synaptonemal complex during meiosis in Caenorhabditis elegans. Genetics. 2011 189 (2): 411–21.

16. Schmekel K, Daneholt B. The central region of the synaptonemal complex revealed in three dimensions. Trends Cell Biol. 1995 5(6): 239–42.

17. Schucker K, Holm T, Franke C, Sauer M, Benavente R. Elucidation of synaptonemal complex organization by super-resolution imaging with isotropic resolution. Proceedings of the National Academy of Sciences of the United States of America. 2015 112(7): 2029–33.

18. von D, Rasmussen SW, Holm PB. The synaptonemal complex in genetic segregation. Annual review of genetics. 1984 18: 331–413.

19. Fraune J, Brochier-Armanet C, Alsheimer M, Volff JN, Schucker K, Benavente R. Evolutionary history of the mammalian synaptonemal complex. Chromosoma. 2016 125 (3): 355–60

20. de Vries FA, de Boer E, van den Bosch M, Baarends WM, Ooms M, Yuan L, et al. Mouse Sycp1 functions in synaptonemal complex assembly, meiotic recombination, and XY body formation. Genes & development. 2005 19 (11): 1376–89.

21. Higgins JD, Sanchez-Moran E, Armstrong SJ, Jones GH, Franklin FC. The Arabidopsis synaptonemal complex protein ZYP1 is required for chromosome synapsis and normal fidelity of crossing over. Genes & development. 2005 19 (20): 2488–500.

22. MacQueen AJ, Colaiacovo MP, McDonald K, Villeneuve AM. Synapsis-dependent and - independent mechanisms stabilize homolog pairing during meiotic prophase in C. elegans. Genes & development. 2002 16 (18) 2428–42.

23. Sym M, Roeder GS. Crossover interference is abolished in the absence of a synaptonemal complex protein. Cell. 1994 79 (2): 283–92.

24. Voelkel-Meiman K, Cheng SY, Morehouse SJ, MacQueen AJ. Synaptonemal Complex Proteins of Budding Yeast Define Reciprocal Roles in MutSgamma-Mediated Crossover Formation. Genetics. 2016 203 (3): 1091–103.

25. Voelkel-Meiman K, Johnston C, Thappeta Y, Subramanian VV, Hochwagen A, MacQueen AJ. Separable Crossover-Promoting and Crossover-Constraining Aspects of Zip1 Activity during Budding Yeast Meiosis. PLoS genetics. 2015 11 (6): e1005335.

26. Colaiacovo MP, MacQueen AJ, Martinez-Perez E, McDonald K, Adamo A, La Volpe A, et al. Synaptonemal complex assembly in C. elegans is dispensable for loading strand-exchange proteins but critical for proper completion of recombination. Developmental cell. 2003 5 (3): 463–74.

27. Smolikov S, Eizinger A, Hurlburt A, Rogers E, Villeneuve AM, Colaiacovo MP. Synapsis-defective mutants reveal a correlation between chromosome conformation and the mode of double-strand break repair during Caenorhabditis elegans meiosis. Genetics. 2007 176 (4): 2027–33

28. Smolikov S, Schild-Prufert K, Colaiacovo MP. A yeast two-hybrid screen for SYP-3 interactors identifies SYP-4, a component required for synaptonemal complex assembly and chiasma formation in Caenorhabditis elegans meiosis. PLoS genetics. 2009 5 (10): e1000669.

29. Hayashi M, Mlynarczyk-Evans S, Villeneuve AM. The synaptonemal complex shapes the crossover landscape through cooperative assembly, crossover promotion and crossover inhibition during Caenorhabditis elegans meiosis. Genetics. 2010 186 (1): 45–58.

30. Libuda DE, Uzawa S, Meyer BJ, Villeneuve AM. Meiotic chromosome structures constrain and respond to designation of crossover sites. Nature. 2013 502 (7473): 703–6.

31. Dernburg AF, McDonald K, Moulder G, Barstead R, Dresser M, Villeneuve AM. Meiotic recombination in C. elegans initiates by a conserved mechanism and is dispensable for homologous chromosome synapsis. Cell. 1998 94 (3): 387–98.

32. Preibisch S, Saalfeld S, Tomancak P. Globally optimal stitching of tiled 3D microscopic image acquisitions. Bioinformatics. 2009 25 (11): 1463–5.

33. Rosu S, Zawadzki KA, Stamper EL, Libuda DE, Reese AL, Dernburg AF, et al. The C. elegans DSB-2 protein reveals a regulatory network that controls competence for meiotic DSB formation and promotes crossover assurance. PLoS genetics. 2013 9 (8): e1003674.

34. Stamper EL, Rodenbusch SE, Rosu S, Ahringer J, Villeneuve AM, Dernburg AF. Identification of DSB-1, a protein required for initiation of meiotic recombination in Caenorhabditis elegans, illuminates a crossover assurance checkpoint. PLoS genetics. 2013 9 (8): e1003679.

35. Woglar A, Daryabeigi A, Adamo A, Habacher C, Machacek T, La Volpe A, et al. Matefin/SUN-1 phosphorylation is part of a surveillance mechanism to coordinate chromosome synapsis and recombination with meiotic progression and chromosome movement. PLoS genetics. 2013 9 (3): e1003335.

36. Jaramillo-Lambert A, Ellefson M, Villeneuve AM, Engebrecht J. Differential timing of S phases, X chromosome replication, and meiotic prophase in the C. elegans germ line. Dev Biol. 2007 308 (1): 206–21.

37. Kelly WG, Schaner CE, Dernburg AF, Lee MH, Kim SK, Villeneuve AM, et al. X-chromosome silencing in the germline of C. elegans. Development. 2002 129 (2): 479–92.

38. Harper NC, Rillo R, Jover-Gil S, Assaf ZJ, Bhalla N, Dernburg AF. Pairing centers recruit a Polo-like kinase to orchestrate meiotic chromosome dynamics in C. elegans. Developmental cell. 2011 21 (5): 934–47.

39. Labella S, Woglar A, Jantsch V, Zetka M. Polo kinases establish links between meiotic chromosomes and cytoskeletal forces essential for homolog pairing. Developmental cell. 2011 21 (5): 948–58.

40. Moses MJ, Poorman PA. Synaptosomal complex analysis of mouse chromosomal rearrangements. II. Synaptic adjustment in a tandem duplication. Chromosoma. 1981 81 (4): 519–35

41. Henzel JV, Nabeshima K, Schvarzstein M, Turner BE, Villeneuve AM, Hillers KJ. An asymmetric chromosome pair undergoes synaptic adjustment and crossover redistribution during Caenorhabditis elegans meiosis: implications for sex chromosome evolution. Genetics. 2011 187 (3): 685–99.

42. MacQueen AJ, Phillips CM, Bhalla N, Weiser P, Villeneuve AM, Dernburg AF. Chromosome sites play dual roles to establish homologous synapsis during meiosis in C. elegans. Cell. 2005 123 (6): 1037–50.

43. Voelkel-Meiman K, Moustafa SS, Lefrancois P, Villeneuve AM, MacQueen AJ. Full-length synaptonemal complex grows continuously during meiotic prophase in budding yeast. PLoS genetics. 2012 8 (10): e1002993.

44. Roelens B, Schvarzstein M, Villeneuve AM. Manipulation of Karyotype in Caenorhabditis elegans Reveals Multiple Inputs Driving Pairwise Chromosome Synapsis During Meiosis. Genetics. 2015 201 (4): 1363–79.

45. Rosu S, Libuda DE, Villeneuve AM. Robust crossover assurance and regulated interhomolog access maintain meiotic crossover number. Science. 2011 334 (6060): 1286–9.

46. Sourirajan A, Lichten M. Polo-like kinase Cdc5 drives exit from pachytene during budding yeast meiosis. Genes & development. 2008 22 (19): 2627–32.

47. Machovina TS, Mainpal R, Daryabeigi A, McGovern O, Paouneskou D, Labella S, et al. Surveillance System Ensures Crossover Formation in C. elegans. Current biology: CB. 2016.

48. Villeneuve AM, Hillers KJ. Whence meiosis? Cell. 2001 106 (6): 647–50.

49. Brenner S. The genetics of Caenorhabditis elegans. Genetics. 1974 77 (1): 71–94.

50. Paix A, Folkmann A, Rasoloson D, Seydoux G. High Efficiency, Homology-Directed Genome Editing in Caenorhabditis elegans Using CRISPR-Cas9 Ribonucleoprotein Complexes. Genetics. 2015 201 (1): 47–54.

51. Martinez-Perez E, Villeneuve AM. HTP-1-dependent constraints coordinate homolog pairing and synapsis and promote chiasma formation during C. elegans meiosis. Genes & development. 2005 19 (22): 2727–43.

52. Schindelin J, Arganda-Carreras I, Frise E, Kaynig V, Longair M, Pietzsch T, et al. Fiji: an open-source platform for biological-image analysis. Nature methods. 2012 9 (7): 676–82.

53. Loidl J, Nairz K, Klein F. Meiotic chromosome synapsis in a haploid yeast. Chromosoma. 1991 100 (4): 221–8.

54. Zhang W, Miley N, Zastrow MS, MacQueen AJ, Sato A, Nabeshima K, et al. HAL-2 promotes homologous pairing during Caenorhabditis elegans meiosis by antagonizing inhibitory effects of synaptonemal complex precursors. PLoS genetics. 2012 8 (8): e1002880.

55. Rog O, Dernburg AF. Direct Visualization Reveals Kinetics of Meiotic Chromosome Synapsis. Cell reports. 2015.

